# Asymmetric framework motion of TCR*αβ* controls load-dependent peptide discrimination

**DOI:** 10.1101/2023.09.10.557064

**Authors:** Ana C. Chang-Gonzalez, Robert J. Mallis, Matthew J. Lang, Ellis L. Reinherz, Wonmuk Hwang

## Abstract

Mechanical force is critical for the interaction between an *αβ*T cell receptor (TCR) and a peptide-bound major histocompatibility complex (pMHC) molecule to initiate productive T-cell activation. However, the underlying mechanism remains unclear. We use all-atom molecular dynamics simulations to examine the A6 TCR bound to HLA-A*02:01 presenting agonist or antagonist peptides under different extensions to simulate the effects of applied load on the complex, elucidating their divergent biological responses. We found that TCR *α* and *β* chains move asymmetrically, which impacts the interface with pMHC, in particular the peptide-sensing CDR3 loops. For the wild-type agonist, the complex stabilizes in a load-dependent manner while antagonists destabilize it. Simulations of the C*β* FG-loop deletion, which reduces the catch bond response, and simulations with *in silico* mutant peptides further support the observed behaviors. The present results highlight the combined role of interdomain motion, fluctuating forces, and interfacial contacts in determining the mechanical response and fine peptide discrimination by a TCR, thereby resolving the conundrum of nearly identical crystal structures of TCR*αβ*-pMHC agonist and antagonist complexes.

## Introduction

The *αβ* TCR (*αβ*TCR) consists of the heterodimeric receptor TCR*αβ* formed by *α* and *β* chains each containing the pMHC-binding variable (V) and constant (C) domains (***Figure 1***A), and the noncovalently associated cluster of differentiation 3 (CD3) subunits that have cytoplasmic tails containing motifs for downstream signaling (***Rudolph, et al. 2006***; Wang and Reinherz, 2012; ***Brazin et al., 2018***). The TCR recognizes its cognate pMHC on the surface of an antigen presenting cell at low or even single copy numbers from a pool of about 10^5^ different self-pMHC molecules (***Sykulev et al., 1996***; ***Brameshuber et al., 2018***), while it also exhibits reactivity with certain closely related peptide variants, with similar or strikingly altered functional T-cell responses (***Ding et al., 1999***; ***Hausmann et al., 1999***; ***Lee et al., 2004***; ***Borbulevych et al., 2009***; ***Baker et al., 2012***; ***Birnbaum et al., 2014***). Considering the *μ*M to hundreds of *μ*M TCR*αβ*-pMHC equilibrium binding affinity (Wang and Reinherz, 2012), several models have been proposed to account for the exquisite specificity and sensitivity of the *αβ*TCR (***Chakraborty and Weiss, 2014***; ***Brazin et al., 2015***; ***Schamel et al., 2019***; ***Zhu et al., 2019***; ***Mariuzza et al., 2020***; ***Liu et al., 2021***).

**Figure 1.**
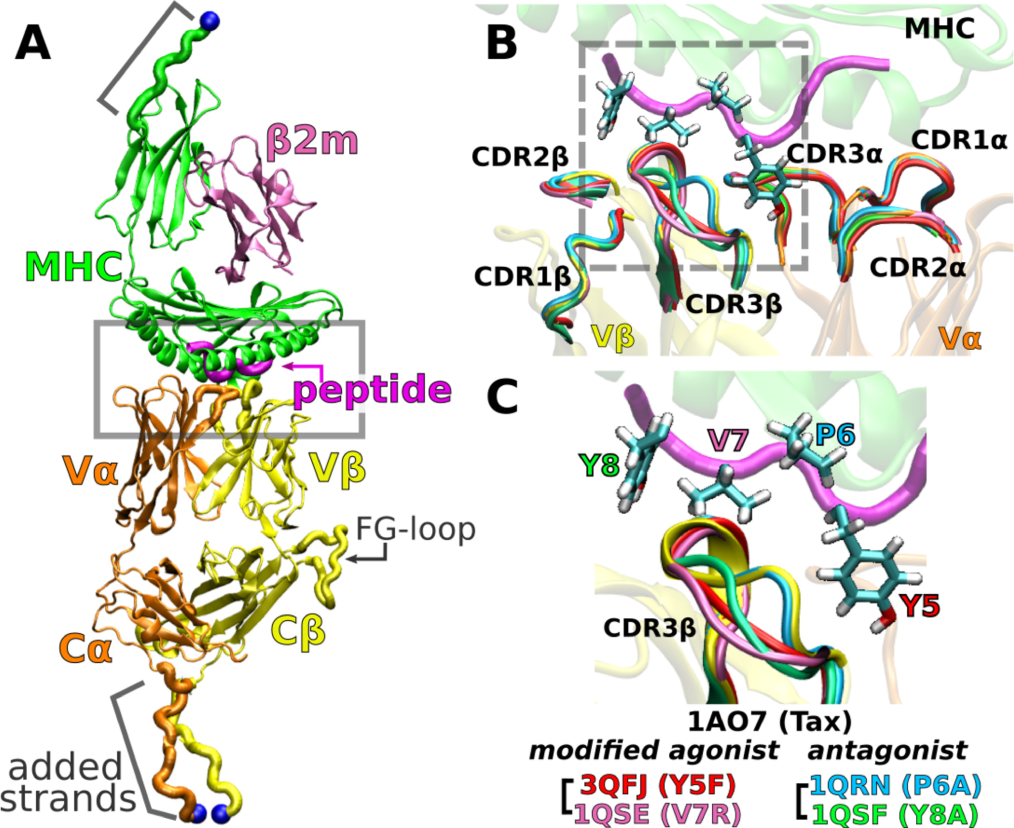
A6 TCR*αβ*-pMHC complex. (A) WT (Protein Data Bank, PDB 1AO7). The missing C*α* domain was added based on PDB 1QSE (Structure preparation). Blue spheres: terminal C_*α*_ atoms held at set extensions during the simulation (***Table 1***). *β*2m: *β*2 microglobulin. (B) Overlay of the X-ray structures of the WT and four point mutants of the Tax peptide at the boxed region of panel A. The CDR loops take nearly identical conformations in different structures. (C) Magnified view of the dashed box in panel B, focusing on the conformation of CDR3*β*. PDB names for A6 TCR*αβ*-pMHC complexes containing mutant peptides are listed.

A critical factor for peptide discrimination is physiological force applied to the TCR*αβ*-pMHC complex (***Reinherz et al., 2023***). A cognate peptide antigen elicits a catch bond behavior where the TCR*αβ*-pMHC bond lifetime increases with force that peaks in the 10–20 pN range, and is observed with the clonotypic ligand-binding TCR*αβ* heterodimer in isolation or with the holoreceptor *αβ*TCR including the non-covalently associated CD3 signaling subunit dimers (CD3*ϵγ*, CD3*ϵγ*, and CD3*ϵδ*). The catch bond is coupled with a roughly 10-nm structural transition in both (***Das et al., 2015***, ***2016***; ***Banik et al., 2021***), which supports the notion that the *αβ*TCR acts as a mechanosensor (***Kim et al., 2009***, ***2012***; ***Brazin et al., 2015***, ***2018***; ***Choi et al., 2023***; ***Reinherz et al., 2023***). In our previous molecular dynamics (MD) study (***Hwang et al., 2020***), instead of enforcing dissociation of the complex with high force, as done in steered MD simulations (***Sibener et al., 2018***; ***Wu et al., 2019***), we applied pN-level forces and examined the behavior of the JM22 TCR complexed with an HLA-A*02:01 molecule presenting a peptide from an influenza virus matrix protein. We found that the TCR*αβ*-pMHC complex is in a loosely-bound state in the absence of load, which allows domain motion. Application of a 16-pN force suppresses the motion and overall enhances the fit among domains. We proposed a model where the TCR*αβ*-pMHC catch bond arises due to stabilization of the interface by altering the conformational motion of TCR*αβ*.

An important question regards the generality of this dynamic mechanism in other TCRs. To this end, we study the A6 TCR, which recognizes the Tax peptide (LLFGYPVYV) of the human T lymphotropic virus type 1 (Garboczi et al., 1996b) bound to HLA-A*02:01, the same MHC as for JM22. We perform all-atom MD simulations with the Tax peptide (wild type; WT) (Garboczi et al., 1996a) and four mutant peptides with a single-residue substitution: Y5F (***Scott et al., 2011***), V7R, P6A, and Y8A (***Ding et al., 1999***). Below, we call the TCR*αβ*-pMHC complex by the name of the corresponding peptide. For example, Y5F refers to the complex with the Y5F peptide (PDB 3QFJ, ***Figure 1***C).

While the crystallographic structures of these complexes are very similar (***Ding et al., 1999***; ***Scott et al., 2011***) (***Figure 1***C), they differ in immunogenicity. P6A and Y8A are weak antagonists because they inhibit T-cell function only at 1000-times higher molar concentration than that needed by the WT for activation (***Hausmann et al., 1999***; ***Ding et al., 1999***). We refer to them simply as “antagonists.” Y5F is similar to WT in terms of equilibrium binding affinity and T-cell activation *in vitro* (***Hausmann et al., 1999***; ***Scott et al., 2011***). V7R induces effector functions comparable to WT at 10- to 100-times higher concentrations (***Ding et al., 1999***). We call Y5F and V7R as “modified ago-nists.” There have been several experimental and computational studies comparing the effects of peptide modifications or pMHC binding on A6 TCR (***Baker et al., 2000***; ***Michielin and Karplus, 2002***; ***Davis-Harrison et al., 2005***; ***Cuendet and Michielin, 2008***; ***Borbulevych et al., 2009***; ***Cuendet et al., 2011***; ***Scott et al., 2011***; ***Ayres et al., 2016***; ***Fodor et al., 2018***), but load was not explicitly considered. To simulate a complex under load, we held the distance between the terminal C_*α*_ atoms of the complex (***Figure 1***A) at a set extension for the duration of the simulation. This was done by applying harmonic positional restraints so that the terminal C_*α*_ atoms were allowed to fluctuate in position, hence resulting in instantaneous fluctuation in the applied force akin to loading in experiments. We refer to a simulation as either low or high load based on the average load, which was around the physiological 10–20 pN range (***Table 1***). To our knowledge, the present study is the first to elucidate the dynamic mechanism of the A6 complex harboring different peptides under load.

We found that differences between the WT and peptide mutants lie in dynamic responses to applied load. In the WT, physiological level load stabilized the TCR*αβ*-pMHC interface as well as the subdomain motion within TCR*αβ*. Modified agonists maintained stable contacts, yet high loads led to destabilization. Antagonists had less stable interfaces under load as the mutated residues disrupted surrounding interfacial contacts. Motion within the TCR, such as the V*α*-V*β* scissoring as observed in ***Hwang et al. (2020)***, and an asymmetric bending of the V-module relative to the C-module, were coupled to the interactions between the variable domains and pMHC such that a single-residue mutation in the peptide affected the conformational behavior of the whole TCR. The present results suggest that the conserved TCR*αβ* framework motion is leveraged when determining the mechanically matched pMHC, a mechanism that is broadly applicable to different TCR*αβ* systems.

## Results

We first study the WT-based systems to gain insight into the functional implications of TCR*αβ*-pMHC structural dynamics, followed by point mutations in the Tax peptide. Our analyses below involve time-dependent inter-domain contact dynamics, domain motion, and their dependence on applied load.

### Load stabilizes WT TCR*αβ*-pMHC interfacial contacts

We assessed the effect of load on WT first by counting high-occupancy contacts with pMHC (Figure 2A; Contact analysis). WT^low^ had the least number of contacts, followed by WT^0^ and WT^high^, indicating low and high loads may have opposite effect on the interfacial stability. V*αβ*-pMHC without the C-module formed the most contacts. This indicates that without a proper load, the C-module is detrimental to the stability of the interface with pMHC, as we found previously for the JM22 TCR (***Hwang et al., 2020***).

Time-dependent changes in the interfacial contacts were monitored using the Hamming distance  (***Hamming, 1950***; ***Hwang et al., 2020***).  is the number of the initial high-occupancy contacts (those with greater than 80% average occupancy during the first 50 ns) that are subsequently lost during the simulation. A low  means that such contacts persist while a high  means the corresponding number of initial high-occupancy contacts are missing. Consistent with the contact count,  remained low for WT^high^ and V*αβ*-pMHC (***Figure 2***B). In WT^0^ and WT^low^,  increased after about 500 ns and 900 ns, respectively. Thus, the relatively high number of interfacial contacts for WT^0^ (***Figure 2***A) is due to the formation of new contacts rather than by maintaining the initial contacts.

**Figure 2.**
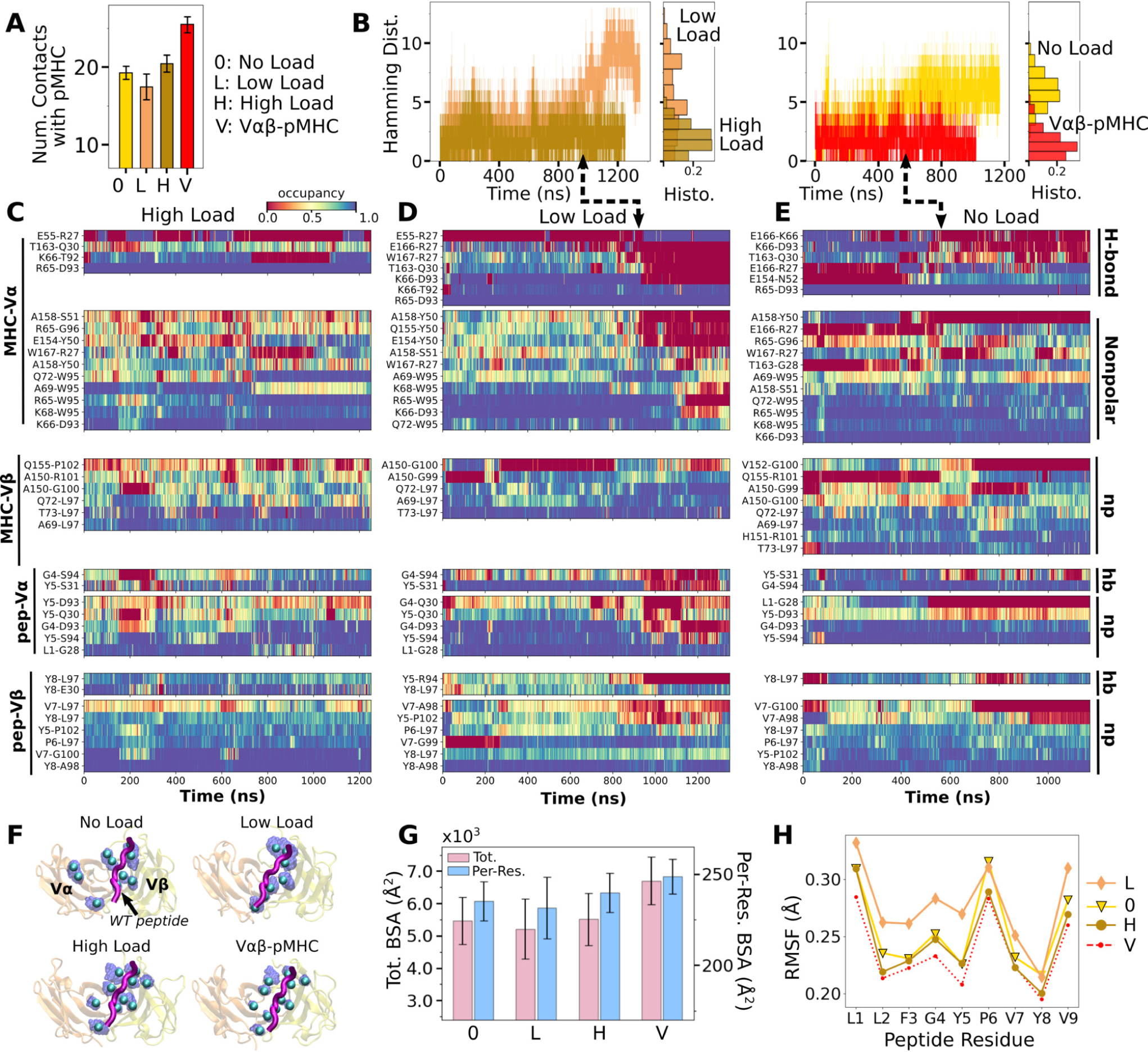
Load dependence of the WT TCR*αβ*-pMHC interface. (A) Number of high-occupancy contacts (Contact analysis). Bars: std. (B) Hamming distance  over time. Histograms are for the interval after 500 ns. Dashed arrows mark increase in , corresponding to contacts lost. (C–E) Contact occupancy heat maps for the interface with pMHC. H-bond/hb: hydrogen bonds, including salt bridges, and np: nonpolar (Contact analysis). (F) Location of C_*α*_ atoms of the residues whose contacts with pMHC have greater than 80% average occupancy. Cyan spheres: last frame of simulation. Transparent blue: locations rendered every 0.2 ns showing positional variability. (G) Total (pink) and per-residue (blue) BSA for interfacial residues with greater than 80% maximum instantaneous occupancy (BSA). Bars: std. (H) RMSF of backbone C_*α*_ atoms of the peptide after 500 ns. Figure 2—figure supplement 1. Contact occupancy heat maps for V*αβ*-pMHC.

Occupancy heat maps provide the time dependence of individual contacts. For WT^high^ and V*αβ*pMHC, high-occupancy contacts persist throughout the simulation (blue regions in ***Figure 2***C and ***Figure 2—figure Supplement 1***) while WT^0^ or WT^low^ exhibited breakage of contacts, especially when  increased (dashed arrows in ***Figure 2***D,E). Differences in the interfacial contacts also manifest in their location. We displayed the backbone C_*α*_ atoms of V*α* and V*β* residues that form contacts with pMHC with greater than 80% average occupancy (averaging was done after the initial 500 ns; ***Figure 2***F). In WT^0^, contacts are spread apart, and in WT^low^ they lie mostly along the length of the peptide. These layouts potentially make interfacial contacts more prone to break via easier access by water molecules. In WT^high^ and V*αβ*-pMHC, high-occupancy contacts form more compact clusters. Exposure to water of the TCR*αβ* residues involved in those contacts was measured by their buried surface area (BSA), which follows the same trend as the number of high occupancy contacts (***Figure 2***A vs. G). Furthermore, this trend also applied to the root-mean square fluctuation (RMSF) of Tax peptide backbone C_*α*_ atoms, even though RMSF values were small (***Figure 2***H).

Experimentally, the WT complex has a relatively strong affinity as a TCR*αβ* (about 1 *μ*M) (***Ding et al., 1999***), which may be why the interface with pMHC involved more contacts in WT^0^ compared to WT^low^. At low load, the short distance between restraints on the ends of the complex (***Figure 1***A) allows wider transverse motion that in turn generates a shear stress or a bending moment at the interface. Such a transverse stress will be less for WT^0^ where the end moves freely, and for WT^high^ where lateral motion is suppressed. The high stability of V*αβ*-pMHC and WT^high^ agree well with the results for JM22 (***Hwang et al., 2020***).

### Influence of pMHC and load on V*α*-V*β* motion

We analyzed the motion between V*α* and V*β* (V*α*-V*β* motion) to find its effect on the TCR*αβ*-pMHC interface. Compared to the unliganded systems (V*αβ* and T*αβ*; ***Table 2***), the number of high-occupancy V*α*-V*β* contacts increased slightly in V*αβ*-pMHC (‘V’ in ***Figure 3***A), while it decreased in full TCR*αβ*pMHC complexes (’0’, ’Low’, and ‘High’ in ***Figure 3***A). This shows that the V*α*-V*β* interface is diffcult to organize with the restrictions imposed by the bound pMHC, except in the absence of the constant domains. The number of V*α*-V*β* contacts in the liganded systems is the smallest for WT^low^, similar to the case for the number of contacts with pMHC (***Figure 2***A), again reflecting a destabilizing effect with low load.

**Figure 3.**
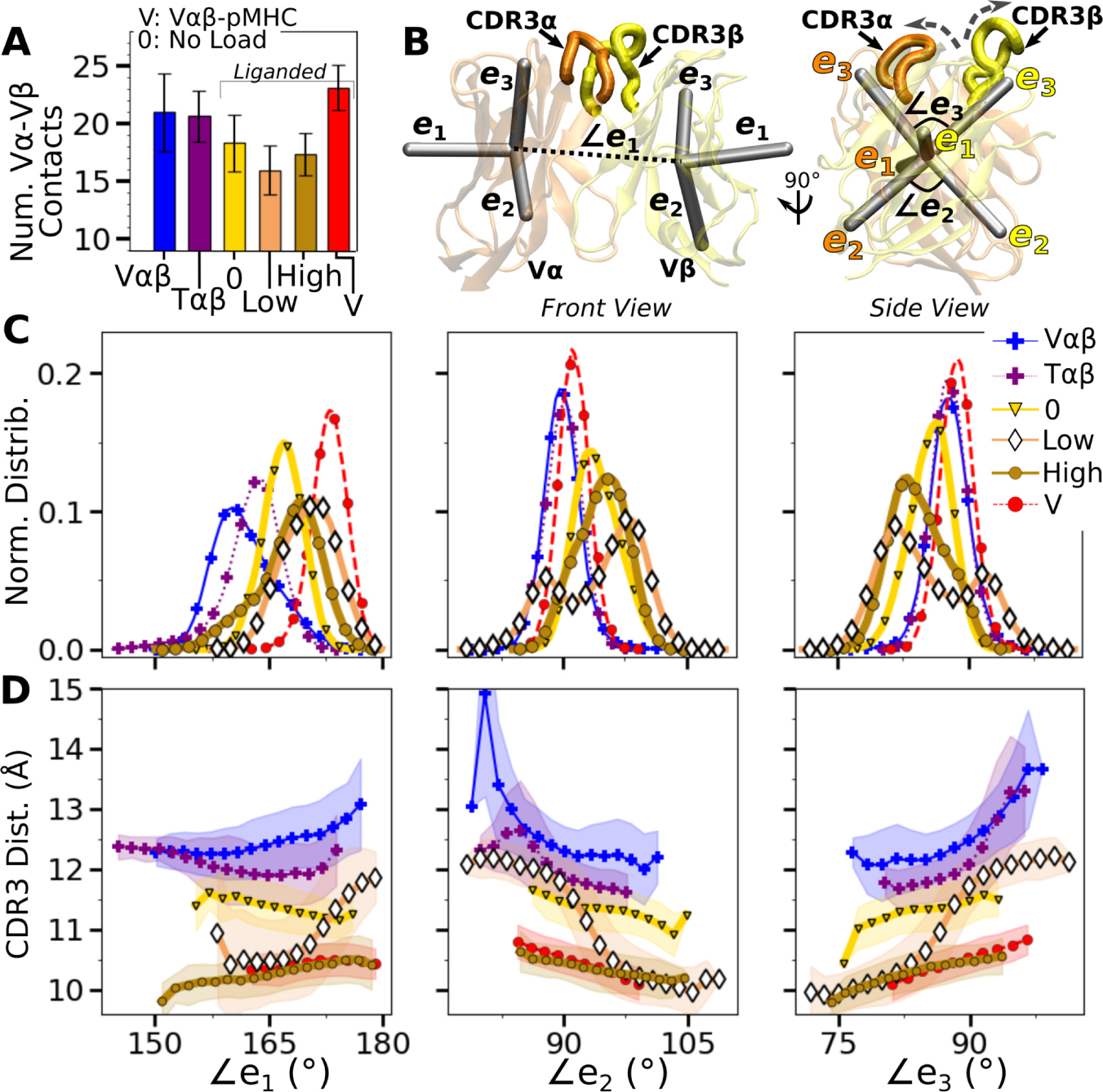
V*α*-V*β* motion. (A) Number of high-occupancy contacts (Contact analysis). Bars: std. (B) Triads {*e*_1_, *e*_2_, *e*_3_} assigned to each domain. Angles between triad arms (∠*e*_1_, ∠*e*_2_, and ∠*e*_3_) are marked. CDR3 loops are represented as thick tubes. Dashed arrows indicate directions where the CDR3 distance increases via the scissor motion. (C) Histograms of the 3 angles between the triad arms. For WT^low^, the smaller peaks in distributions of ∠*e*_2_ and ∠*e*_3_ arise from simulation trajectories after 1 *μ*s. (D) CDR3 distance vs. the 3 angles. Transparent band: std of the CDR3 distance in each bin. Statistics for bins deteriorate in large- or small-angle tails that contain very few frames. Figure 3—figure supplement 1. PCA of V*α*-V*β* motion. Figure 3—figure supplement 2. Trajectories of the V-module motion.

V*α*-V*β* motion was measured by assigning triads to the stably folded *β*-sheet cores of the two domains and performing principal component analysis (PCA) on the trajectories of the two triads (***Figure 3***B; Variable domain triads and PCA) (***Hwang et al., 2020***). The amplitude of PC1 is lower when the number of V*α*-V*β* contacts is higher (***Figure 3***A vs. ***Figure 3—figure Supplement 1A***). Directions of PCs differed to varying extents (arrows in ***Figure 3—figure Supplement 1B***). Similarity of the directions was measured by the absolute value of the dot product between PCs as 18-dimensional unit vectors (for the 6 arms from two triads) in different systems. A value of 1 corresponds to the same direction, and 0 means an orthogonal direction (***Figure 3—figure Supplement 1C***). For PC1, a high degree of similarity was observed between T*αβ* and V*αβ*, which is consistent with their similarity in the number of V*α*-V*β* contacts (***Figure 3***A) and PC amplitudes (***Figure 3—figure Supplement 1A***). Among triad systems with bound pMHC, WT^low^ differed significantly in the PC1 direction compared to others (***Figure 3—figure Supplement 1C***, darker colors). The dot products varied more for PC2 and PC3, which capture finer motions with smaller amplitudes (***Figure 3—figure Supplement 1C***).

To determine how the V*α*-V*β* motion influences the interface with pMHC, we measured the distance between CDR3 loops (CDR3 distance), which play a central role in peptide discrimination (***Figure 1***B,C and ***Figure 3—figure Supplement 2***A–C). Unliganded T*αβ* and V*αβ* had greater fluctuation in the CDR3 distance (larger std), as they are unrestrained by pMHC. Among the pMHC-bound systems, WT^high^ and V*αβ*-pMHC had small CDR3 distance (averages of 10.3 and 10.5 Å, respectively; ***Figure 3—figure Supplement 2***B,C). CDR3 distance was larger for WT^0^ (11.3 Å), which reflects an altered interface with pMHC. For WT^low^, the CDR3 distance varied more widely, with more than a 2-fold increase in standard deviation. The increase in CDR3 distance of WT^low^ happens after the in-crease in  (800 ns; ***Figure 2***B,D and ***Figure 3—figure Supplement 2***C), suggesting a loss of contacts at the interface is related to the V*α*-V*β* motion.

PCA decomposes the V*α*-V*β* motion into mutually orthogonal directions. We made 2-dimensional histograms of each of these projections versus the corresponding CDR3 distance (***Figure 3—figure Supplement 1***D). If any of the PC modes is strongly correlated with the CDR3 distance, the corresponding histogram would exhibit a slanted profile. However, no clear correlation could be seen (***Figure 3—figure Supplement 1***D), suggesting that the changes in CDR3 distance may depend on combinations of PCs. We addressed this possibility by considering angles between matching arms of the two triads (***Figure 3***B). The *e*_1_-*e*_1_ angle, herein called ∠*e*_1_ (and similarly define ∠*e*_2_ and ∠*e*_3_; ***Figure 3***B), can change either by the *e*_1_ arms turning up and down (“flap”) or in and out of the page (“twist”) in ***Figure 3***B. Angles ∠*e*_2_ and ∠*e*_3_ depend primarily on rotation indicated by dashed arrows in ***Figure 3***B (“scissor”) (***Hwang et al., 2020***).

Histograms of the three angles (***Figure 3***C) show a clearer difference than individual PCs among the systems tested, and the CDR3 distance varies with the angles (***Figure 3***D). The wider distributions for angles of WT^low^ and WT^high^ reflect their higher PC amplitudes (***Figure 3—figure Supplement 1***A). The symmetric distributions of ∠*e*_2_ and ∠*e*_3_ can be seen from the two peaks for WT^low^, which is due to the reciprocal behavior of the scissoring motion involving the two angles (***Figure 3***B, side view, and ***Figure 3***C, open diamonds). The two peaks are also related to the changes in the CDR3 distance (***Figure 3***D), which reflects an agitating effect of the mild load on the scissor motion. Given the definitions of the angles, the CDR3 distance will increase (dashed arrows in ***Figure 3***B) with larger ∠*e*_1_ or ∠*e*_3_, or with smaller ∠*e*_2_ (***Figure 3***D). WT^0^, despite the apparent stability of the interface, had a larger CDR3 distance than WT^high^ and V*αβ*-pMHC, again indicating a disrupted interface. These results show that the CDR3 motion is coupled to the V*α*-V*β* motion, especially the scissoring motion.

### Asymmetric V-C motion influences the load response of the complex

We next analyzed the motion of the V-module relative to the C-module (V-C motion). The number of high-occupancy C*α*-C*β* contacts did not vary significantly (in the 31–34 range) and they were more than the number of V*α*-V*β* contacts, similar to the case for the JM22 TCR (***Hwang et al., 2020***). The C-module thereby influences the V-module as a single unit. The V-C motion was analyzed by performing PCA on the bead-on-chain (BOC) model constructed based on the *β*-sheet core of each domain, and hinges between Vand C-domains denoted as H*α* and H*β* (***Figure 4***A; V-C BOC and PCA). Across different systems, amplitudes of PCs were similar (***Figure 4—figure Supplement 1***A). PC1 (V-C bend; ***Figure 4***A) was similar among systems, as seen by the values of dot products being close to 1.0 (***Figure 4***B). Directions of higher PCs varied more, similarly as higher PCs for the V*α*-V*β* motion.

**Figure 4.**
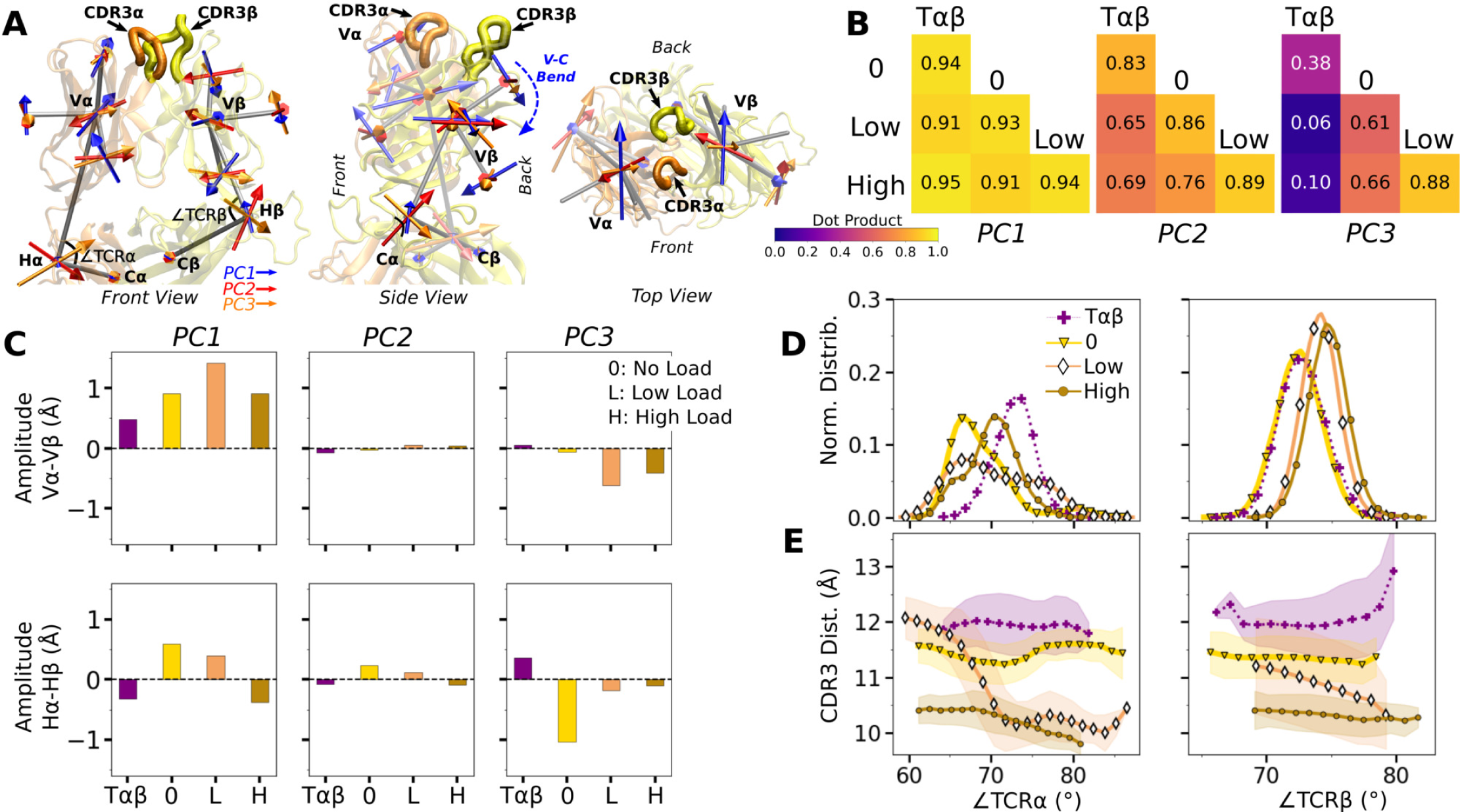
WT V-C dynamics. (A) Average BOC built from the unliganded T*αβ*. (B) Dot products computed between the BOC PCs in listed systems. Values closer to 1.0 denote similar V-C BOC direction of motion. (C) Difference in amplitude between the *α* and *β* chain motion measured between V*α* and V*β* (top), and H*α* and H*β* (bottom). PC amplitudes are proportional to the lengths of the arrows in panel A. (D) Histograms of hinge angles (defined in panel A) for each chain. (E) CDR3 distance vs. hinge angles. Figure 4—figure supplement 1. V-C PC amplitude and contacts.

We noticed that V*α* bends more compared to V*β*, as can be gleaned from the longer PC arrows for V*α* (***Figure 4***A). We quantified this asymmetry by subtracting the amplitudes of motion for domains in the *β* chain from those for the matching domains in the *α* chains, where positive or negative values respectively indicate greater or less motion of the *α* compared to the *β* chain (***Figure 4***C). Compared to T*αβ*, binding of pMHC increases the *α*-chain motion, which is the greatest in WT^low^ (***Figure 4***C, PC1 in top row). The greater degree of V*α*-C*α* motion is consistent with the smaller number of V*α*-C*α* contacts compared to V*β*-C*β* (***Figure 4—figure Supplement 1***B,C).

The asymmetry was further analyzed by measuring hinge angles ∠TCR*α* and ∠TCR*β* (***Figure 4***A). Distributions of ∠TCR*α* varied more compared to ∠TCR*β* (***Figure 4***D). A wide distribution of ∠TCR*α* for WT^low^ is related to the increase in the CDR3 distance and concomitant changes in the triad arm angles later during the simulation (***Figure 3—figure Supplement 2***C). For WT^low^ and WT^high^, CDR3 distance decreases with increasing hinge angles, especially with ∠TCR*α* (***Figure 4***E), which suggests that unbending of the V-module under load helps with bringing the CDR3 loops closer together. In WT^low^, this state is not maintained and ∠TCR*α* decreases (more bending) as the CDR3 distance increases (***Figure 4***E) which happens after the increase in  (***Figure 2***B). These results suggest a mechanism by which the asymmetric response of the whole TCR*αβ* to load affects the binding with pMHC by controlling the relative positioning between the CDR3 loops via the V*α*-V*β* motion. For this, the C*β* FG-loop plays a critical role, as simulations of the bound complex without the C*β* FG-loop resulted in a smaller ∠TCR*β* and an over-extended ∠TCR*α* (see Appendix 1).

### Effects of point mutations on the peptide

In the WT crystal structure, the side chain of Y5 in the Tax peptide is located between the CDR3 loops of V*α* and V*β* while V7 mainly contacts CDR3*β* (***Figure 1***C). P6 makes one contact with CDR3*β*. The side chain of Y8 is located between CDR3*β* and the *α*1 helix of MHC. Crystal structures of point mutants of these four residues are very similar in terms of interfacial contacts, docking angle, and CDR loop conformations, with the only structurally observable difference located at CDR3*β* (***Figure 1***B,C) (***Ding et al., 1999***; ***Scott et al., 2011***). However, point mutations profoundly affect dynamics of the complex, as explained below.

Modified agonists Y5F and V7R had about the same number of contacts with pMHC as the WT complexes, but high load resulted in fewer contacts, indicating a potential slip bond behavior (***Figure 5***A), though loss of contacts in Y5F^high^ might have been due to a higher load experienced compared to other complexes at the same extension (23.7 pN; ***Table 1***, Selecting extensions). Antagonists P6A and Y8A had overall fewer contacts with pMHC without a consistent load dependence (***Figure 5***A). This trend was also seen in BSA profiles of residues forming high-occupancy contacts with pMHC (***Figure 5***B). For modified agonists, higher load also resulted in greater increase of , whereas the trend was opposite for antagonists (***Figure 5—figure Supplement 1***). The large number of contacts with pMHC for modified agonists (***Figure 5***A) despite an increase in  suggests an altered binding rather than maintaining the initial contacts.

**Figure 5.**
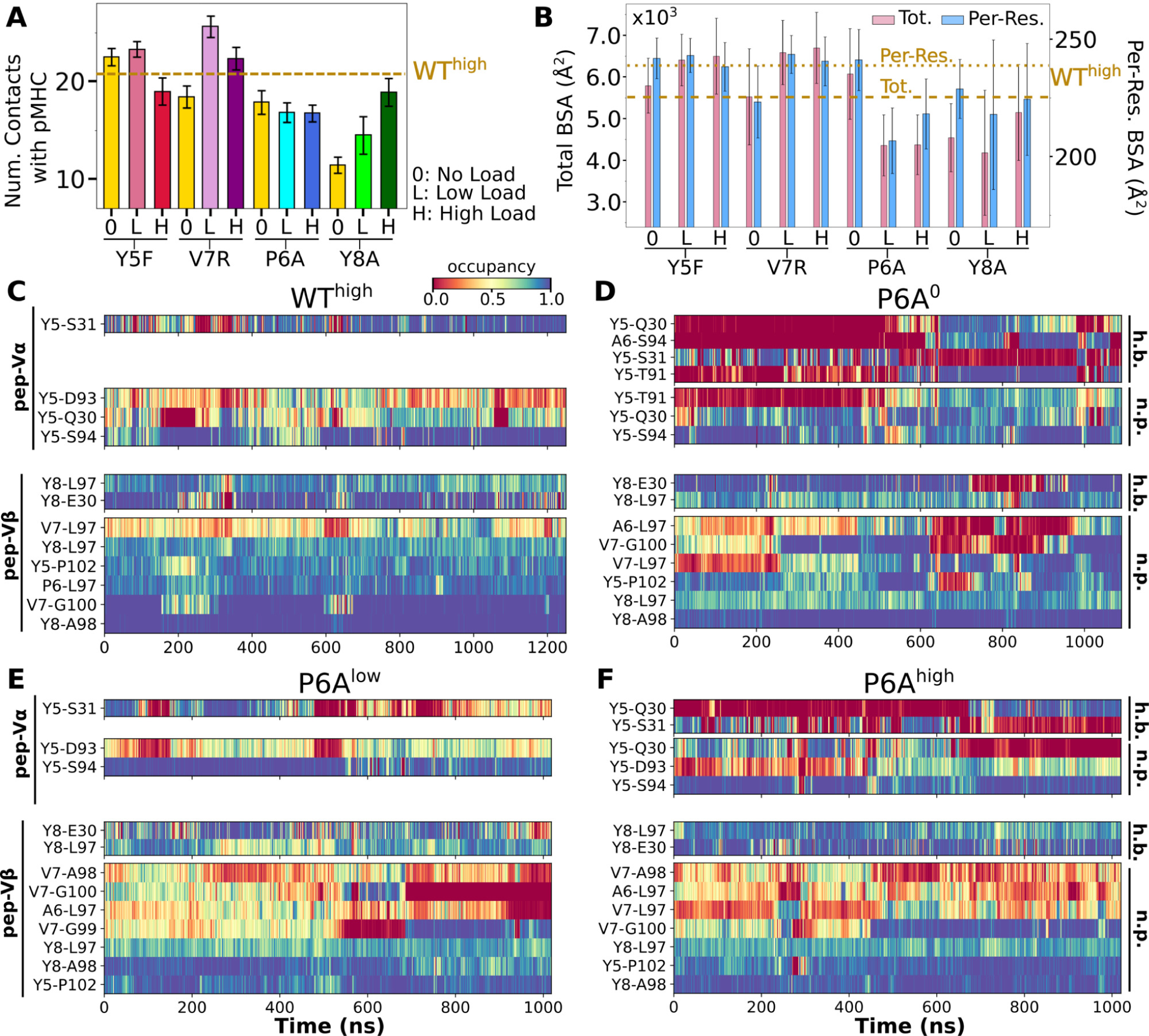
Interface with pMHC containing mutant peptides. The same occupancy cutoffs were used as in ***Figure 2***. (A) Number of contacts with pMHC. Dashed line: count for WT^high^ in ***Figure 2***A, for reference. (B) Total (pink) and per-residue (blue) BSA. Dashed and dotted lines: values for WT^high^ (***Figure 2***G). (C-F) Contact heat maps for peptide residues 5 to 8. (C) WT^high^ (included in ***Figure 2***C), and (D) P6A^0^, (E) P6A^low^, and (F) P6A^high^. Figure 5—figure supplement 1. Trajectories of  for mutant complexes. Figure 5—figure supplement 2. Contact occupancy heat maps for residues 5–8 of Y5F, V7R, and Y8A. Figure 5—figure supplement 3. Locations of high-occupancy contacts with pMHC in mutant systems.

In contact heat maps, the Y5 residue of the WT peptide forms a hydrogen bond with *α*S31 and nonpolar contacts with a few residues in both V*α* and V*β* (***Figure 2***C–E, ***Figure 5***C). In Y5F, the hydrogen bond with *α*S31 cannot form, and many of the nonpolar contacts with F5 break under load later during the simulation (***Figure 5—figure Supplement 2***A). The breakage coincides with the increase in  (***Figure 5—figure Supplement 1***A). In addition, contacts involving Y8 and V7 also break in Y5F^high^ (***Figure 5—figure Supplement 2***A). Thus, the Y5-*α*S31 hydrogen bond may stabilize the interface with pMHC by arranging other nearby residues to form nonpolar contacts in favorable positions; its absence would make the nonpolar contacts more prone to break under load. The relative stability of Y5F^0^ can also be seen by the similarity in the locations of high-occupancy contact residues between WT and Y5F^0^ (***Figure 2***F vs. ***Figure 5—figure Supplement 3***A). Experimentally, Y5F has kinetic and cytotoxicity profiles similar to WT (***Hausmann et al., 1999***; ***Scott et al., 2011***). Its dependence on load needs further experimental analysis. On the other hand, V7 of the WT peptide forms nonpolar contacts with residues in CDR3*β* (***Figure 2***C–E, ***Figure 5***C). In V7R, nonpolar contacts with CDR3*β* form with reduced occupancy, and contacts involving Y8 are also disrupted (***Figure 5***C vs. ***Figure 5—figure Supplement 2***B).

For antagonists, more contacts broke, which again involve non-mutated residues such as Y5 and V7 (***Figure 5***D–F,***Figure 5—figure Supplement 2***C). The greater number of contacts in Y8A^high^ compared to Y8A^0^ and Y8A^low^ (***Figure 5***A) despite smaller number of contacts involving key peptide residues Y5–A8 (***Figure 5—figure Supplement 2***C) suggests formation of additional contacts with other parts of MHC as a result of an altered interface. Experimentally, binding of the A6 TCR to pMHC containing the P6A or Y8A peptide was not detected *in vitro* (***Ding et al., 1999***). Thus, Y8A in principle could exhibit a catch bond, but forming the complex in the loaded state may be kinetically inaccessible.

The modified agonists had more V*α*-V*β* contacts than WT^high^ while the antagonists had fewer, except for Y8A^high^ (***Figure 6—figure Supplement 1***A). While the amplitude of V*α*-V*β* motion was generally in a range similar to the WT systems (***Figure 3—figure Supplement 1***A vs. ***Figure 6—figure Supplement 1***B), the CDR3 distance was larger for all mutant systems except for P6A, which had a weak dependence on triad angles (***Figure 6***A–B). The angles in turn varied among systems and loading conditions (***Figure 6—figure Supplement 2***). These results suggest that point mutations to the peptide cause alterations in the load-dependence of the interface and the V*α*-V*β* motion.

**Figure 6.**
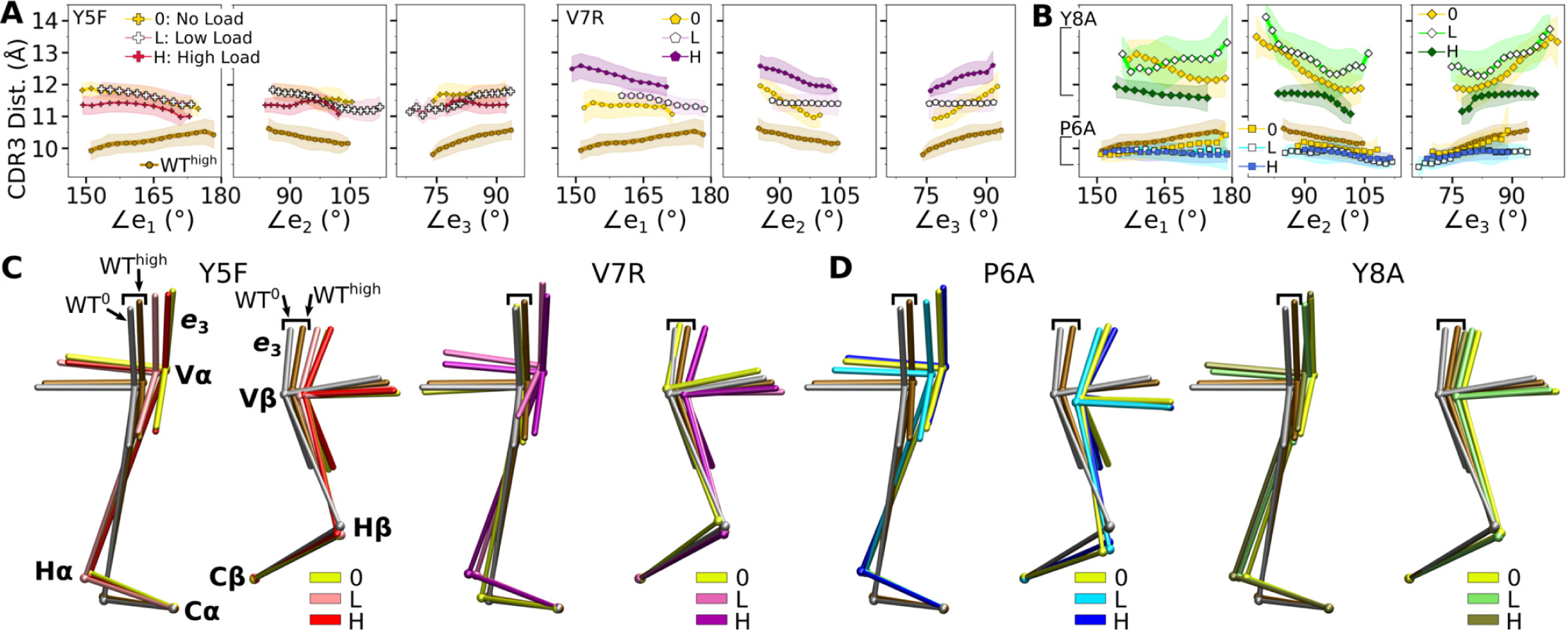
Mutant effects on the conformational dynamics of TCR*αβ*. (A,B) CDR3 distance versus triad arm angles for (A) modified agonists and (B) antagonists. Plot for WT^high^ in ***Figure 3***D is reproduced for comparison. (C,D) Average BOCs of labeled complexes oriented to the constant domains of WT^high^ (V-C BOC and PCA) for (C) modified agonists and (D) antagonists. Average BOCs for WT^0^ and WT^high^ are displayed for comparison (marked by angular brackets). Figure 6—figure supplement 1. V*α*-V*β* motion of mutant systems. Figure 6—figure supplement 2. Distribution of triad arm angles in mutant systems. Figure 6—figure supplement 3. Comparison of mutant average V-C BOCs and interfaces with those of WThigh. Figure 6—figure supplement 4. V-C motion of mutants.

The mutants affected the average V-C BOC similarly as that for dFG^high^ (***Figure 6***C,D vs. Appendix 1–***Figure 1***C). Among them, Y8A^high^ had an average BOC approaching that of WT^high^, which aligns with the comparable number of contacts with pMHC (***Figure 5***A). However, the location of its H*α* differed (***Figure 6***D), and the CDR3 distance was larger (***Figure 6***B). To quantify deformation of the average BOC, we measured displacements of centroids from the corresponding ones in WT^high^. They were overall greater for the *α* chain than the *β* chain (***Figure 6—figure Supplement 3***A,B). Consistent with this, the mutants had fewer V*α*-C*α* contacts than WT^high^ and a similar number of V*β*-C*β* contacts (***Figure 6—figure Supplement 3***C,D).

Similar to the WT systems, the greater motion of the *α* chain than the *β* chain was observed in the mutant systems, as seen from the differences in V-C PC1 amplitudes (***Figure 6—figure Supplement 4***A,B). However, dot products of the BOC PC1 between WT and mutants revealed that the direction of motion differed by varying degrees, which was more for V7R^high^ and Y8A (***Figure 6***— ***figure Supplement 4***C vs. ***Figure 4***B). Thus, point mutations on the WT peptide can affect the conformational motion of the whole TCR*αβ*, in addition to the average BOC.

To further test effects of point mutations, we introduced *in silico* point mutations P6A and Y8A to the WT complex (WT to antagonists) and conversely introduced A6P and A8Y mutations to the P6A and Y8A complexes, respectively (antagonists to WT). The *in silico* antagonists did exhibit reduction in contacts with pMHC while the results were mixed for the *in silico* WT, especially for A8Y where the introduced tyrosine is bulkier than the original alanine. Nevertheless, these tests support the above results based on the original crystal structures (See Appendix 2 for details).

### Load- and time-dependent interfacial response

To probe the dynamic relation between the TCR*αβ*-pMHC (intermolecular) interface and intra-TCR*αβ* (intramolecular) interfaces formed between subdomains of the complex, we calculated the total occupancy of the high-occupancy contacts in respective cases (***Figure 7***A,B). For the intramolecular contacts, we excluded the C*α*-C*β* interface contacts since they are larger in number compared to other interfaces and did not differ significantly across different systems, *i.e.*, the C-module moves mostly as a single unit (***Hwang et al., 2020***).

**Figure 7.**
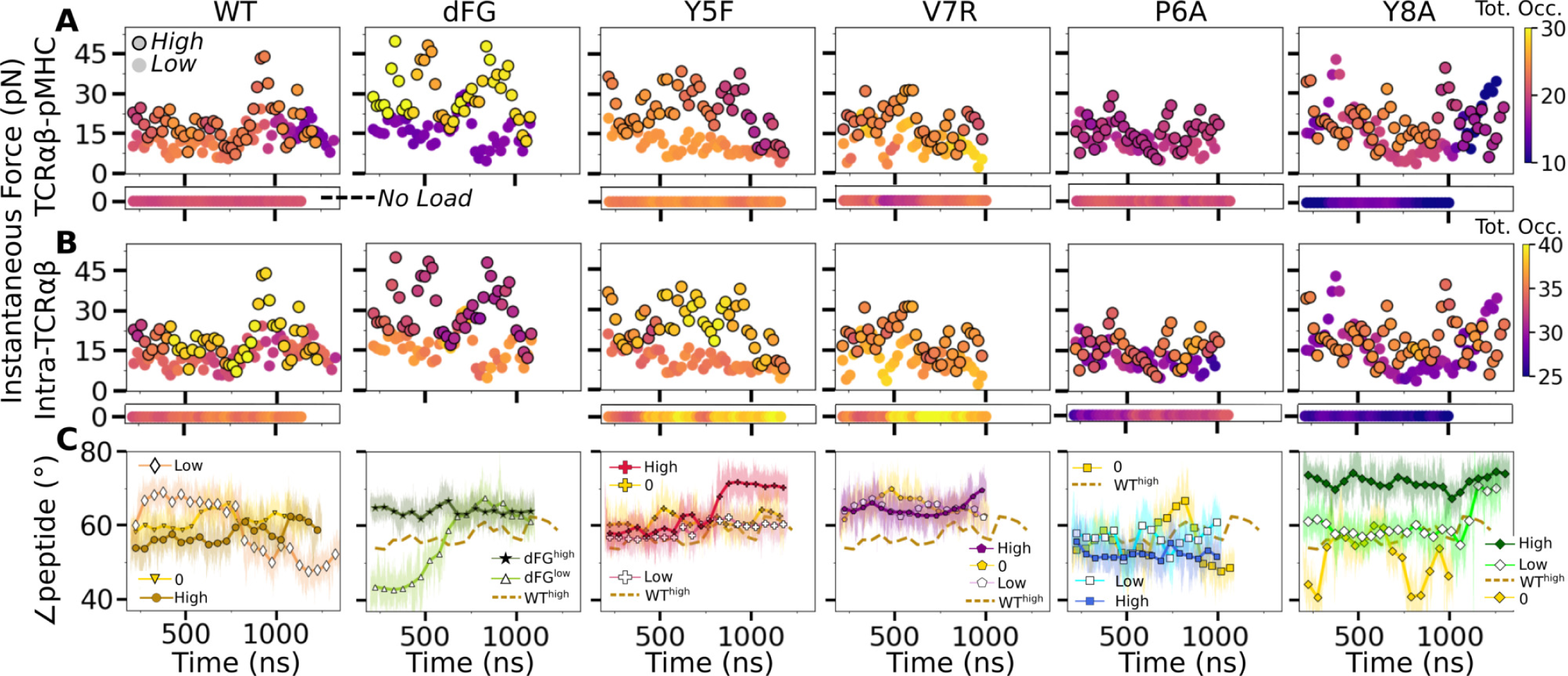
Relationship between force and interfacial behavior. (A,B) The total contact occupancy measured in 40-ns overlapping intervals starting from 200 ns (Time-dependent behavior). (A) TCR*αβ*-pMHC (intermolecular) and (B) intra-TCR*αβ* (intramolecular) contacts excluding C*α*-C*β*. Cases without load are shown as horizontal bars below each panel. Plots for low load systems (Table 1) do not have outlines. (C) Angle between antigenic peptide and the line between centroids of the triads for V*α* and V*β* (Peptide and V-module angle). Thin lines: values at individual frames. Symbol: 50-ns running average. Figure 7—figure supplement 1. Motion at the interface related to ∠peptide.

**Table 1.** Simulations of TCR*αβ*-pMHC complexes. Load is average after 500 ns (See Selecting extensions).

| Peptide | PDB | Extension (Å) | Time (ns) | Load (pN) | Label | Description |
| --- | --- | --- | --- | --- | --- | --- |
| Tax<br>(WT) | 1AO7 | – | 1170 | – | WT<sup>0</sup> | wild-type |
|  |  | 182.6 | 1350 | 13.2 | WT<sup>low</sup> |  |
|  |  | 187.7 | 1250 | 18.2 | WT<sup>high</sup> |  |
| Tax<br>(dFG-pMHC) | 1AO7 | 180.5 | 1100 | 14.9 | dFG<sup>low</sup> | dFG (*Table 2*)<br>with pMHC |
|  |  | 188.9 | 1100 | 29.0 | dFG<sup>high</sup> |  |
| Y5F | 3QFJ | – | 1180 | – | Y5F<sup>0</sup> | modified<br>agonists |
|  |  | 181.4 | 1200 | 8.24 | Y5F<sup>low</sup> |  |
|  |  | 186.2 | 1200 | 23.7 | Y5F<sup>high</sup> |  |
| V7R | 1QSE | – | 1020 | – | V7R<sup>0</sup> | modified<br>agonists |
|  |  | 177.5 | 1012 | 10.3 | V7R<sup>low</sup> |  |
|  |  | 186.2 | 1003 | 17.8 | V7R<sup>high</sup> |  |
| P6A | 1QRN | – | 1090 | – | P6A<sup>0</sup> | weak<br>antagonists |
|  |  | 175.2 | 1018 | 8.81 | P6A<sup>low</sup> |  |
|  |  | 186.0 | 1020 | 13.5 | P6A<sup>high</sup> |  |
| Y8A | 1QSF | – | 1020 | – | Y8A<sup>0</sup> | weak<br>antagonists |
|  |  | 176.5 | 1280 | 12.0 | Y8A<sup>low</sup> |  |
|  |  | 187.4 | 1330 | 18.1 | Y8A<sup>high</sup> |  |

**Table 2.** Simulations of truncated structures from PDB 1AO7.

| Label | Modification | Time (ns) |
| --- | --- | --- |
| Vαβ | Vα-Vβ only (no pMHC) | 1060 |
| Tαβ | TCRαβ only (no pMHC) | 1000 |
| Vαβ-pMHC | Vαβ with pMHC (no C-module) | 1020 |
| dFG | Tαβ without the Cβ FG-loop (no pMHC) | 1000 |

For WT^0^, the intermolecular contact occupancy stayed at around 20 (WT in ***Figure 7***A, horizontal bar on the bottom) and for WT^low^, it decreased later in simulation (WT in ***Figure 7***A, darkening of circles without outline). In comparison, the intramolecular contact occupancy remained relatively constant for both WT^0^ and WT^low^ (WT in ***Figure 7***B, horizontal bar on the bottom and circles without outline). For WT^high^, the intermolecular contact occupancy was steady even with wider fluctuation in force (WT in ***Figure 7***A, outlined circles), and the intramolecular occupancy also remained high, indicating the subdomains are held together tightly (WT in ***Figure 7***B, outlined circles). For dFG^low^, the intermolecular contact occupancy stayed low and intramolecular occupancy was relatively constant (dFG in ***Figure 7***A,B, circles without outline). In dFG^high^, the contact occupancy with pMHC increased (dFG in ***Figure 7***A, outlined circles), but the intramolecular contact occupancy became low (dFG in ***Figure 7***B, outlined circles), which suggests that the complex is not as tightly coupled compared to WT.

For modified agonists, the no load and low load cases had overall higher occupancy, both with pMHC and within TCR*αβ*, but occupancy fluctuated more as can be seen by the changes in colors in the occupancy trajectories (Y5F and V7R in ***Figure 7***A,B, horizontal bars on the bottom and circles without outlines). Under high load, intermolecular contact occupancy decreased over time (Y5F and V7R in ***Figure 7***A, darkening of outlined circles) while intramolecular contact occupancy either increased (Y5F^high^) or decreased (V7R^high^) relative to the respective low load cases. For antagonists, both occupancy measures were lower than the WT, and further reduction could be seen over time in some cases (P6A and Y8A in ***Figure 7***A,B, darkening of colors in outlined circles).

The stability of the TCR*αβ*-pMHC interface also manifested into their relative motion, which was quantified by the angle between the least-square fit line across the backbone C_*α*_ atoms of the antigenic peptide and the unit vector formed between the centroids of V*α* and V*β* (***Figure 7***— ***figure Supplement 1***A–D). For WT, the peptide angle fluctuated more for WT^low^ than WT^high^ (WT in ***Figure 7***C) where 58.4^◦^±3.6^◦^ (avg±std after 500 ns) for WT^high^ reflects a diagonal binding. For dFG^low^, the peptide changed orientation by more than 20^◦^, and for dFG^high^, it stabilized, but at a higher value than WT^high^, which also was reached in dFG^low^ later during simulation, suggesting a more orthogonal binding (dFG in ***Figure 7***C). For modified agonists, similar to the behaviors of the total intra- and intermolecular contact occupancy, the peptide angle was affected more under high loads, again becoming more orthogonal compared to WT^high^ (Y5F and V7R in ***Figure 7***C). For antagonists, the angle overall fluctuated more under no load or settled to different values under high load. Since the antagonists are loosely coupled (low occupancy in ***Figure 7***A,B), settling of the angle does not indicate stabilization of the interface, as evident from the positional shift of the *α*2 helix of V7R^high^ or Y8A^high^ (***Figure 7—figure Supplement 1***G,H) compared to WT^high^ (***Figure 7—figure Supplement 1***E).

## Concluding Discussion

The present study elucidates how the load-dependent TCR*αβ* framework motion influences the dynamics of the TCR*αβ*-pMHC interface (***Figure 8***). A main feature of the framework is the smaller number of contacts for the V*α*-C*α* compared to the V*β*-C*β* interface. This causes an asymmetric V-C motion, primarily bending, where V*α* moves more compared to V*β* relative to the C-module, which serves as a base. This in turn generates relative motion between V*α* and V*β*, which can destabilize the contacts with pMHC, especially by affecting the distance between CDR3 loops that play the most direct role for sensing the bound peptide (***Figure 8***A,B). Applying a physiological level force stabilizes the interface by straining the whole complex into a more tightly coupled state, as can be seen by the increase of both inter- and intramolecular contacts in WT^high^ (***Figure 7*** and ***Figure 8***C). The CDR3 distance of WT^high^ (10.3±0.3 Å; ***Figure 3—figure Supplement 2***C) was shorter than that of WT^0^ or WT^low^ (***Figure 3***D), and it is also shorter than the 10.9-Å CDR3 distance in the crystal structure (PDB 1AO7). The applied load slightly increases the spacing between pMHC and TCR*αβ*, which provides room for the CDR3 loops to adjust as well as allow other contacts to ‘lock’ into more stable states with higher and more persistent occupancy. Absence of load or low load do not properly channel the framework motion and thereby increase exposure to water (***Figure 2***F,G) and destabilize the interface.

**Figure 8.**
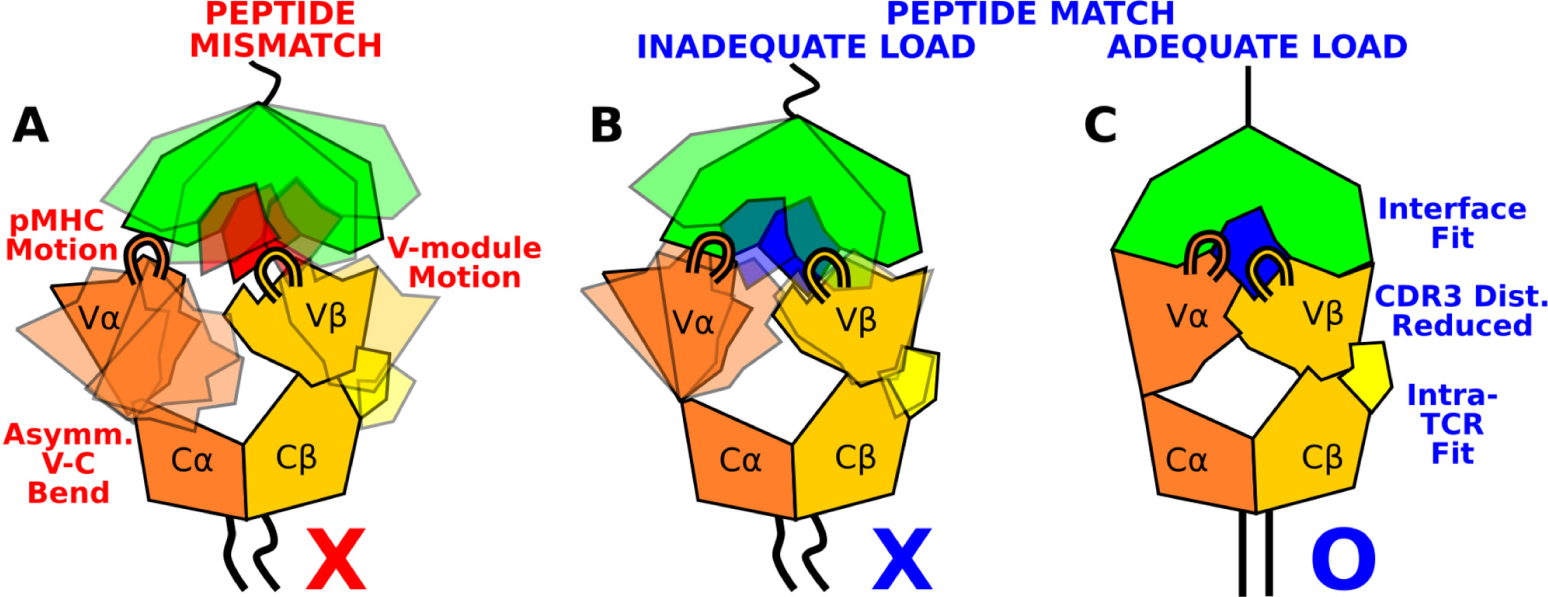
Model for peptide screening. (A) Non-matching pMHC or (B) matching pMHC but without adequate load do not stabilize the asymmetric V-C framework motion that affects the interfacial stability as measured by the CDR3 distance (CDR3 loops are shown above the V-module). (C) Matching pMHC with adequate load results in an overall tighter fit.

The C*β* FG-loop stabilizes the V*β*-C*β* interface, thereby contributing to the asymmetric V-C motion. It also controls the relative orientation between V*β* and C*β*, hence it affects the orientation of the CDR loops of the V-module with respect to the loading direction (Appendix 1–***Figure 1***C). Consistency in these findings between the present study and our previous simulations using JM22 TCR (***Hwang et al., 2020***) underscores that the proposed mechanism based on the asymmetric framework motion is applicable to other TCR*αβ* systems.

After engagement with a cognate pMHC under load, reversible transition to an extended state is possible which has been observed both in *in vitro* single-molecule experiment using TCR*αβ* and on cell displaying the full *αβ*TCR holoreceptor (***Das et al., 2015***; ***Banik et al., 2021***). Since V*αβ*-pMHC lacking the C-module forms a more stable binding (***Figure 2***) that was also observed in our previous simulations of the JM22 TCR (***Hwang et al., 2020***), the C-module likely undergoes partial unfolding in the extended state. Thus, while the folded C-module serves as the base for the asymmetric V-C motion screening for the matching pMHC, once a match is found, the reversible transitioning propelled by the partial unfolding of the C-module may agitate the membrane and activate the cytoplasmic domains of the surrounding CD3 subunits to initiate downstream signaling (***Reinherz et al., 2023***). A circumstantial evidence for the capacity of the C-module to unfold is that the C*α* domain as well as parts of C*β* are occasionally unresolved in crystal structures, as in PDB 1AO7.

In addition to TCR*αβ*, MHC may also respond to load. ***Wu et al. (2019)*** suggested a partial separation of the MHC*α*1-*α*2 peptide-binding platform from *β*2m with the attendant lengthening of pMHC contributing to a longer bond lifetime. ***Banik et al. (2021)*** observed a catch bond for CAR-pMHC, where just MHC is being pulled with an antibody. While we did not find a clear load or peptide-dependence in contacts between subdomains of MHC, since the entire TCR*αβ*-pMHC complex is under load, conformational changes in pMHC may contribute to the extended state of the complex. Yet, for T-cell based cancer immunotherapy, mechanistic knowledge of the mechanosensing through a TCR has a greater practical significance (***Reinherz et al., 2023***).

A recent study using a laminar flow chamber assay fit the measured bead survival distribution using Bell’s equation to estimate the zero-force off rate *K*_off_ and the force sensitivity distance *x*_*β*_ (***Pettmann et al., 2022***). They found a negative correlation between *K*_off_ and *x*_*β*_, to conclude that mechanical forces impair antigen discrimination. However, the force range tested was up to 100 pN, where even systems exhibiting catch bond in the 10–20-pN range will switch to a slip bond behavior.

A catch bond exhibits a non-monotonic force versus bond lifetime profile, so that fitting with Bell’s equation, an exponential function, leads to results that do not have a clear physical meaning. For example, *x*_*β*_ in ***Pettmann et al. (2022)*** was less than 1 Å in magnitude in all systems, which is shorter than the length of a single covalent bond. They also performed steered MD simulation that applies hundreds of pN forces, which is inadequate for studying behaviors of the system under loads in the 10–20-pN range (***Hwang et al., 2020***). Use of a coarse grained model without appropriately incorporating atomistic properties of the TCR further makes it difficult to compare their simulation with experiment.

We earlier proposed that the residues of the antigenic peptide play a role more as “teeth of a key” for screening the TCR*αβ*-pMHC interaction fitness rather than bearing applied loads (***Hwang et al., 2020***; ***Reinherz et al., 2023***). The present study confirms this through simulations of mutant systems, where several contacts across the interface with pMHC were impaired due to a single-residue mutation on the peptide in ways that reflect the functional outcome of the mutation. In considering how a T-cell may respond to an unknown peptide, the pMHC motion and the asymmetric V-C motion are two points of guidance (***Figure 8***). Stabilization of the inter- and intramolecular interfaces throughout the whole complex under 10–20-pN load would indicate a cognate TCR*αβ*pMHC interaction (***Figure 7***). Since these features are based on overall TCR*αβ*-pMHC complex dynamics, rather than changes to specific contacts or a particular conformational change, they can be used to predict fitness of other TCR*αβ*-pMHC combinations. Since such tests involve performing many all-atom MD simulations and trajectory analyses, an *in silico* method would be needed that effciently predicts dynamic properties of the complex based on sequence and structural data only. Atomistic insights gained from the present study will be helpful for developing such a method in future studies.

## Computational Methods

### Structure preparation

Structure preparation was done using CHARMM (***Brooks et al., 2009***). Simulation systems were based on PDB 1AO7 (Garboczi et al., 1996a); 1QSE, 1QRN, and 1QSF (***Ding et al., 1999***); and 3QFJ (***Scott et al., 2011***) (***Figure 1***C). Residues from the TCR *α*- and *β*-chains were renumbered sequentially from the original non-sequential numbering in the PDB. Throughout the paper we use the renumbered index to refer to a residue. Residues differing at a few locations in some of the PDB files were converted so that all systems have identical sequences except for point mutations introduced in the Tax peptide (details are given below). Disulfide bonds between cysteine residues were introduced as noted in the PDB file. Histidine protonation sites were determined based on the 1QSE crystal structure to promote hydrogen bond formation with neighboring residues. Where neighboring residues were unlikely to hydrogen bond, we assigned the water-facing nitrogen of histidine as charged. This led to protonation of the N^*γ*^ atom for all histidine residues except for MHC H263 and *β*2m H84, where the N^*ϵ*^ atom was protonated. For truncated structures, crystal waters within 2.8 Å from the protein atoms were kept in the initially built system. For full structures, all crystal waters were kept.

We extended the termini of the TCR*αβ*-pMHC complex as handles for applying positional restraints (***Figure 1***, “added strands”) (***Hwang et al., 2020***). For MHC, we used the sequence from UniProt P01892, where ^276^LSSQPTIPI^284^ was added after E275. For TCR*αβ*, sequences for the added strands were from GenBank ABB89050.1 (TCR*α*) and AAC08953.1 (TCR*β*), which were ^201^PESSCDVK LVEKSFETDT^218^ and ^246^CGFTSESYQQGVLSA^260^, respectively. After adding the strands, a series of energy minimization and a short MD simulation in the FACTS implicit solvent environment (Haberthür and Caflisch, 2008) were performed to relax them and bring together the C-terminal ends of the two TCR chains. The first two N-terminal residues of TCR*β* were missing in all structures except for 3QFJ, so they were added and briefly energy minimized.

### 1AO7 (Tax peptide)

In the original PDB 1AO7, coordinates for the C*α* domain (D116–S204) and parts of C*β* (E130–T143, K179–R188, S219–R228) are missing. The coordinates listed are based on the renumbered indices. These were built using PDB 1QSE. For the C*α* domain, we aligned the V*α* domain of 1AO7 and 1QSE (K1–P115) based on their backbone C_*α*_ atoms and added the missing C*α* domain residue coordinates to 1AO7. After this, we performed a brief energy minimization on the added domain while fixing positions of all other atoms of 1AO7. For missing residues in the C*β* domain, we used backbone C_*α*_ atoms of two residues each before and after the missing part to align 1QSE to 1AO7 and filled in coordinates, followed by a brief energy minimization of the added part in 1AO7. In this way, the TCR*αβ*-pMHC interface of the original 1AO7 is preserved. By comparison, previous simulations mutated PDB 1QRN back to WT (***Ayres et al., 2016***), which corresponds to to the A6P *in silico* WT system (Appendix 2), or converted a high-affinity variant of A6 (PDB 4FTV) by mutating *β*-chain residues, in particular nearly the entire CDR3 loop (***Rangarajan et al., 2018***). Compared to our approach, those preparation methods thereby introduce more perturbation to the interface with pMHC.

The *β*2m residues C67 and C91 were reverted (C67Y, C91K) based on UniProt P61769 referenced in PDB 1AO7. These agree with the *β*2m sequence in other structures.

### 1QRN (P6A)

Except for the two N-terminal residues of TCR*β*, there were no missing coordinates.

This also applies to 1QSE and 1QSF. The following conversions were made to match the sequence with other structures: K150S (TCR*α*), and A133E and E134A (TCR*β*).

1QSE (V7R): No residue conversion was made.

1QSF (Y8A): The following conversions were made: A219R (MHC) and A225T (TCR*β*).

3QFJ (Y5F): There were no missing residues. We made the D204N conversion in TCR*β*.

WT truncated complexes: For truncation, we used the constructed 1AO7 complex.

- V*αβ*: the last residues were *α*D111 and *β*E116.
- T*αβ*: the last residues were *α*D206 and *β*G247 (no C-terminal strands).
- V*αβ*-pMHC: includes V*αβ*, peptide, *β*2m, and MHC. The last residue of MHC was L276.
- WT^0^: WT complex without the added C-terminal strands, as for T*αβ*.
- dFG: residues *β*L218–*β*P231 removed from the corresponding WT complex. *β*G217 and *β*V232 were covalently joined.

### MD simulation protocol

Solvation and equilibration of simulated systems We used CHARMM (***Brooks et al., 2009***) to prepare simulation systems before the production run. The solvation boxes were orthorhombic for systems with pMHC and cubic for those without pMHC. For TCR*αβ*-pMHC, the size of the initial water box was such that protein atoms were at least 12 Å away from the nearest transverse face of the box and 25 Å from each longitudinal face. The extra space in the longitudinal direction was to initially test and select extensions of the complex for longer simulations in ***Table 1***. For solvation, we used the TIP3P water. Water molecules with their oxygen atoms less than 2.8 Å from protein heavy atoms were removed. Neutralization of the system was done using Na^+^ and Cl^−^ ions at about 50 mM concentration. Crystal water molecules were kept in this procedure.

After neutralization, a 5-stage energy minimization was applied where protein backbone and side chain heavy atoms were progressively relaxed (***Hwang et al., 2020***). This was followed by heating from 30 K to 300 K for 100 ps and equilibration at 300 K for 200 ps. Backbone heavy atoms were positionally restrained with 5-kcal/[mol⋅Å^2^] harmonic spring constant during heating and equilibration, except for structures involving 1AO7 that originally had more missing residues, where a 2-kcal/[mol⋅Å^2^] restraint was used. We then performed a 2 ns CPT (constant pressure and temperature) simulation at 1 atm and 300 K. We applied a 0.001-kcal/[mol⋅Å^2^] restraint on backbone C_*α*_ atoms. The CHARMM DOMDEC module (***Hynninen and Crowley, 2014***) was used to parallelize the simulation. We applied the SHAKE method to fix the length of covalent bonds involving hydrogen atoms, and used a 2-fs integration time step.

### Production runs

Production runs were performed using OpenMM (***Eastman et al., 2017***). We used the CHARMM param36 all-atom force field (***Huang and MacKerell Jr, 2013***) and the particle-mesh Ewald method to calculate electrostatic interactions. We used an Ewald error tolerance of 10^−5^ which is 1/50 of the default value in OpenMM, for accuracy. The cutoff distance for nonbonded interactions was 12 Å, and the Nose-Hoover integrator of OpenMM at 300 K was used, with a 2-fs integration time step. We ran OpenMM on GPUs with mixed floating point precision. Below are specific steps of the MD protocol relevant to individual systems in ***Table 1*** and ***Table 2***.

### TCR*αβ*-pMHC with load

#### Laddered extension with added strands

To apply load, C_*α*_ atoms of the C-terminal ends of the added strands in the complex (***Figure 1***A, blue spheres) were held by 1-kcal/[mol⋅Å^2^] harmonic positional restraint at a given extension during the simulation. Restraints were applied to the C_*α*_ atom of MHC I284 and to the center of mass of two C_*α*_ atoms of *α*T218 and *β*A260. A flat-bottom distance restraint was applied to the latter two atoms to prevent large separation. It was activated when the distance of the two C_*α*_ atoms was greater than 10 Å, where a 1.0-kcal/[mol⋅Å^2^] harmonic potential was applied. Starting with the initially built complex, we performed a 4-ns run then increased the extension by shifting centers of the positional restraints on terminal atoms by 2 Å at each end, for a total 4 Å added at each extension, for the next 4-ns run. The process continued to yield 4–6 extensions.

After each extension run, we truncated the water box such that the length of the box was 12 Å larger than the maximum span of the complex on each side, and re-neutralized the system. A representative water box size is 218×97×90 Å^3^ for WT^high^, containing 187,250 atoms. Since the system was already equilibrated from the previous run, we used a simpler energy minimization scheme where backbone and side chain heavy atoms were restrained by 10-kcal/[mol⋅Å^2^] and 5-kcal/[mol⋅Å^2^] harmonic potentials, respectively, and 200 steps of steepest descent followed by 200 steps of adopted-basis Newton-Raphson energy minimization was performed. Heating, equilibration, and the initial 2-ns dynamic runs with positional restraints were carried out as explained above. We then carried out 60–100 ns production runs for each extension and selected two or three extensions to continue for longer than 1000 ns.

### Selecting extensions

We measured the average force on the complex during each 60–100-ns simulation, then selected two extensions where the average force generated was representative of a “low” (around 10 pN) and “high” (over 15 pN) load on the TCR. These values were based on the experimental 10–20-pN catch bond activation force range (***Das et al., 2015***; ***Liu et al., 2016***).

In some cases, in particular at low extensions, the flexible added strand either folded onto itself or made contacts with the C-module of TCR*αβ*, effectively shortening the span of the complex. Factors such as this, together with differences in conformational behaviors of the complex, affected the average force for a given extension. Thus we had to test and choose among different extensions for each system. We also ran 1–2 replicate simulations of comparable length (∼1 *μ*s) at given extensions except for systems involving dFG and *in silico* mutants. However, even with nearly the same extensions used, measured forces in replicate simulations varied. The final selection and average forces are in ***Table 1***.

### Other systems

#### TCR*αβ*-pMHC without load

These systems include WT^0^ and complexes with point mutations to the Tax peptide. To prevent the complex from turning transversely in the elongated orthorhombic box, we applied a weak 0.2- kcal/[mol⋅Å^2^] harmonic positional restraint on select C_*α*_ atoms in the MHC *α*3 domain that had RMSF below about 0.5 Å in both WT^low^ and WT^high^, which were P185–T187, L201–Y209, F241–V247, and T258–H263.

#### V*αβ*-pMHC

We applied a 0.01-kcal/[mol⋅Å^2^] harmonic restraint to the backbone C_*α*_ atoms of the MHC *α*3 domain (residues P185-L276) to prevent the complex from turning transversely in the orthorhombic box. The restraints are 20 times weaker than those used for TCR*αβ*-pMHC complexes mentioned above. This was because V*αβ*-pMHC is smaller in both size and aspect ratio.

#### V*αβ*, T*αβ*, dFG

No positional restraints were applied. A representative system size is, for T*αβ*, a 92.8-Å^3^ cubic water box containing 75,615 atoms.

#### dFG-pMHC

The FG-loop deletion was done after initially preparing (solvation and neutralization) the WT complex in the extended water box. After deletion, the system was re-neutralized. Subsequently, laddered extension, selecting extensions for high and low load cases, and longer production runs were performed as explained above.

### In silico mutants

Each *in silico* mutation (Appendix 2–***Table 1***) was performed for low and high load extensions of the complex. To use similar extensions as in the original complexes, we used the last frame of the 4-ns laddered extension simulation. After introducing the *in silico* mutation, we inspected the structure to ensure there was no steric clash with neighboring residues or water molecules. We performed a short energy minimization to relax the modified residue while keeping coordinates of all other residues except for residues immediately before and after the mutated one on the peptide. We then truncated the water box and re-neutralized the system, after which steps from the initial energy minimization up to the final production run followed the same procedure as explained above.

### Trajectory Analysis

Coordinates were saved every 20 ps (0.02 ns) during production runs, resulting in 50,000 coordinate frames for 1000 ns. We excluded the initial 500 ns when calculating averages and standard deviations in the number of contacts, CDR3 distance, BSA, PCA values, and angle data. Since all systems were simulated for a minimum of 1 *μ*s, this leaves at least 25,000 frames. We report data prior to 500 ns in trajectory plots and contact occupancy heat maps (e.g., ***Figure 2***B–E).

### Calculating force

Force on a restrained atom or the center of mass of the C-terminal atoms of the added strands in TCR*αβ* was calculated based on the deviation of its average position from the center of the harmonic potential, multiplied by the spring constant used (***Hwang et al., 2020***). Average force in ***Table 1*** was computed from 500 ns to the end of the simulation. Instantaneous force in ***Figure 7***A,B was computed in 40-ns overlapping intervals starting from 200 ns, *i.e.*, 200–240 ns, 220–260 ns, 240–280 ns, *etc*.

### RMSF

RMSF for backbone C_*α*_ atoms of a domain was calculated by aligning the C_*α*_ atoms to the structure at the beginning of the production run. Coordinate frames after the initial 500 ns were used.

### CDR3 distance

The CDR3 distance (e.g. ***Figure 3—figure Supplement 2***A–C) was measured using the midpoint between backbone C_*α*_ atoms of two residues at the base of each CDR3 loop. They were: T92 and K97 for CDR3*α*, and R94 and E103 for CDR3*β*.

### Contact analysis

We used our previously developed method (***Hwang et al., 2020***). Briefly, H-bonds (including salt bridges) were identified with the 2.4-Å donor-acceptor distance cutoff. Nonpolar contacts were identified for atom pairs that are within 3.0 Å and both have partial charges less than 0.3*e* (*e* = 1.6 × 10^−19^ C) in magnitude. The average occupancy was measured as the fraction of frames over which a bond is present during the measurement period. Instantaneous occupancy was measured as a 40-frame (0.8-ns) rolling average. The average occupancy of a contact represents its abundance during the simulation period while the instantaneous occupancy represents its temporal intensity.

For counting the number of contacts (e.g., ***Figure 2***A, ***Figure 3***A, ***Figure 4—figure Supplement 1***B,C), we used contacts with the average occupancy greater than 50% and at least an 80% maximum instantaneous occupancy after the initial 500 ns. Contact occupancy heat maps (e.g., ***Figure 2***C–E) report those with the overall average occupancy greater than 30%, and the maximum instantaneous occupancy during the simulation greater than 80%.

The Hamming distance  (e.g. ***Figure 2***B) was measured using contacts with greater than 80% average occupancy during the first 50 ns.

### BSA

For the BSA calculation (e.g. ***Figure 2***G), we used residues in the V-module with the maximum instantaneous contact occupancy with pMHC greater than 80%. We calculated the surface area for the selected residue contacts and added them to get the total BSA. Per-residue BSA is the total BSA divided by the number of residues forming the contacts in the given time interval. The reported values (e.g. ***Figure 2***G) are respective averages after 500 ns.

### Variable domain triads and PCA

Triads (orthonormal unit vectors) were constructed for V*α* and V*β* by modifying the procedure in ***Hwang et al. (2020)*** for the A6 V-module. We used the backbone C_*α*_ atoms of six residues from the central four *β*-strands that make up the stably folded *β*-sheet core of each variable domain: for V*α*, S19-Y24, F32-Q37, Y70-I75, Y86-T91, and for V*β*, T20-Q25, S33-D38, F74-L79, V88-S93. The C_*α*_ atoms of these residues have RMSF in WT^high^ near or less than 0.5 Å, and they correspond to two matching segments on each of the inner and outer *β*-sheets of the immunoglobulin fold. The center of mass of the C_*α*_ atoms of the selected residues was used for the centroid of each triad. The *e*_3_ arm of the triad was assigned along the major axis of the least-square fit plane of the selected atoms in each domain, which is parallel to the *β*-strands and points to the CDR3 loop (***Figure 3***B). The *e*_1_ arm was assigned by taking the direction from the center of masses of the selected atoms from the inner to the outer *β*-sheets of each variable domain and making it perpendicular to *e*_3_. The *e*_2_ arm was then determined as *e*_2_ = *e*_3_ × *e*_1_.

PCA was performed on the trajectory of the two triads using a custom FORTRAN95 program (***Hwang et al., 2020***). The PC amplitude (e.g. ***Figure 3—figure Supplement 1***A) corresponds to the rotational motion of these arms in units of radians. The PC vector for the 6 arms of the two triads is an 18-dimensional unit vector. To compare directions of two PCs (e.g. ***Figure 3—figure Supplement 1***C), the absolute value of the dot product between them was calculated, which ranges between 0 and 1. To project the V*α*-V*β* triad for a given frame to a PC direction (***Figure 3—figure Supplement 1***D), the average triad calculated after the initial 500-ns was subtracted from the triad, then a dot product was formed with the PC vector.

### V-C BOC and PCA

The V-C BOC (***Figure 4***A) was assigned based on the method we developed previously (***Hwang et al., 2020***). For beads representing the V-module, centroids of the two triads were used. For the Cmodule, the center of mass of backbone C_*α*_ atoms of the following residues in each domain were used: for C*α*, A118–R123, V132–D137, Y153–T158, S171–S176, and for C*β*, T143–A148, L158–N163, S192–V197, F209–Q214. We used *α*N114 for H*α*, and for H*β*, the center of mass between *β*D117 and *β*L118 was used, which had large RMSF in WT^high^.

We aligned coordinate frames for all simulations to the first frame of WT^high^ based on atoms used to assign beads for the C-module. In this way, motion of the V-module relative to the C-module can be analyzed. Also, by using a common reference structure (first frame of WT^high^), average BOCs can be compared, as in ***Figure 6***C,D. PCA of the V-C BOC was performed using the 6 beads representing the centroids and hinges. PCA for the V-module triads was done separately. Since the reference of motion is the C-module, directions of PCs for the V-module triads indicate motion of the V-module relative to the C-module (arrows on triad arms in ***Figure 4***A), which complements the direction of the V-module centroids obtained from PCA of the V-C BOC (arrows on centroids in ***Figure 4***A).

### Time-dependent behavior

For the total occupancy in ***Figure 7***A,B, we only considered contacts with greater than 50% overall occupancy and over 80% maximum instantaneous occupancy during the entire simulation period. In this way, changes in high-quality contacts under fluctuating force for a given trajectory can be monitored. For each 40-ns window, we calculated the average occupancy of selected contacts and added them to obtain the total occupancy. For intermolecular contacts, interfaces between MHC-V*α*, MHC-V*β*, peptide-V*α*, and peptide-V*β* were considered. For intramolecular contacts, V*α*-V*β*, V*α*-C*α*, and V*β*-C*β* were considered.

### Peptide and V-module angle

For ***Figure 7***C, at each coordinate frame we calculated the least-square fit line for the peptide backbone C_*α*_ atoms and calculated a dot product of its direction with a unit vector pointing from the centroid for the triad of V*α* to that of V*β*.

## Acknowledgments

This work was funded by US NIH Grants P01AI143565 and R01AI136301. Simulations were performed by using computers at the Texas A&M High Performance Research Computing facility.

### Appendix 1

#### Allosteric effect of the C*β* FG-loop deletion

The higher number of V*β*-C*β* contacts (***Figure 4—figure Supplement 1***B,C) and smaller range of ∠TCR*β* (***Figure 4***D,E) in the WT TCR suggests that the *β* chain mainly bears the load while the *α* chain adjusts to accommodate different loading conditions. Our previous experimental (***Das et al., 2015***) and computational (***Hwang et al., 2020***) studies showed that the C*β* FG-loop plays a critical allosteric role for the catch bond formation. To examine its role in A6 TCR, we performed simulations of dFG in isolation (***Table 2***) and under low and high loads (***Table 1***). Similar to the WT, the number of contacts with pMHC increased with load (Appendix 1–***Figure 1***A and Appendix 1–***Figure 2***A,B). The BSA for high-occupancy residues contacting pMHC was also greater for high load (Appendix 1–***Figure 2***C). Thus, dFG may also possess a catch bond behavior, which agrees with experiment where a subdued catch bond was observed (***Das et al., 2015***). However,  increased early on and was slightly larger than that for WT^high^ (Appendix 1–***Figure 2***D), and the CDR3 distance was higher compared to WT^high^ (Appendix 1–***Figure 1***B), which indicate an altered interface.

The conformation of the whole dFG was also affected. Relative to the C-module, the average BOC for the unliganded dFG was substantially different from those of dFG^low^ and dFG^high^, where the latter was similar to that of WT^high^ (Appendix 1–***Figure 1***C). In particular, V*β* of the unliganded dFG is more tilted, as there is a lack of support from the FG-loop (***Hwang et al., 2020***). When load is applied to the dFG-pMHC complex, dFG becomes less bent. Its tendency to return to the bent conformation would impose a strain on the interface with pMHC. This can be seen by the higher average load on dFG-pMHC complexes than WT-pMHC complexes under similar extensions (***Table 1***). Comparing between the amplitudes of PCs of *α* and *β* chains, a notable difference from the WT systems (***Figure 4***C) is that H*β* moves more than H*α* for loaded dFG systems (Appendix 1–***Figure 1***D, bottom row). Also, distributions of ∠TCR*α* and ∠TCR*β* shift to larger and smaller values, respectively (Appendix 1–***Figure 1***E). These indicate alterations in the conformation and motion of dFG. Furthermore, the CDR3 distance of dFG is elevated regardless of load or V-C angle (Appendix 1–***Figure 1***F), suggesting a reduced allosteric control by the V-C motion.

The altered conformation of dFG causes the interface with pMHC to tilt as observed in our previous study of JM22, which is detrimental to the stability of the complex (***Hwang et al., 2020***). The increased motion of H*β* may also deliver more agitation to the interface with pMHC. Thus, even though full dissociation with pMHC was not observed within the simulation time, the dFG-pMHC complex is likely to be less stable compared to the WT complex.

**Appendix 1—figure 1.**
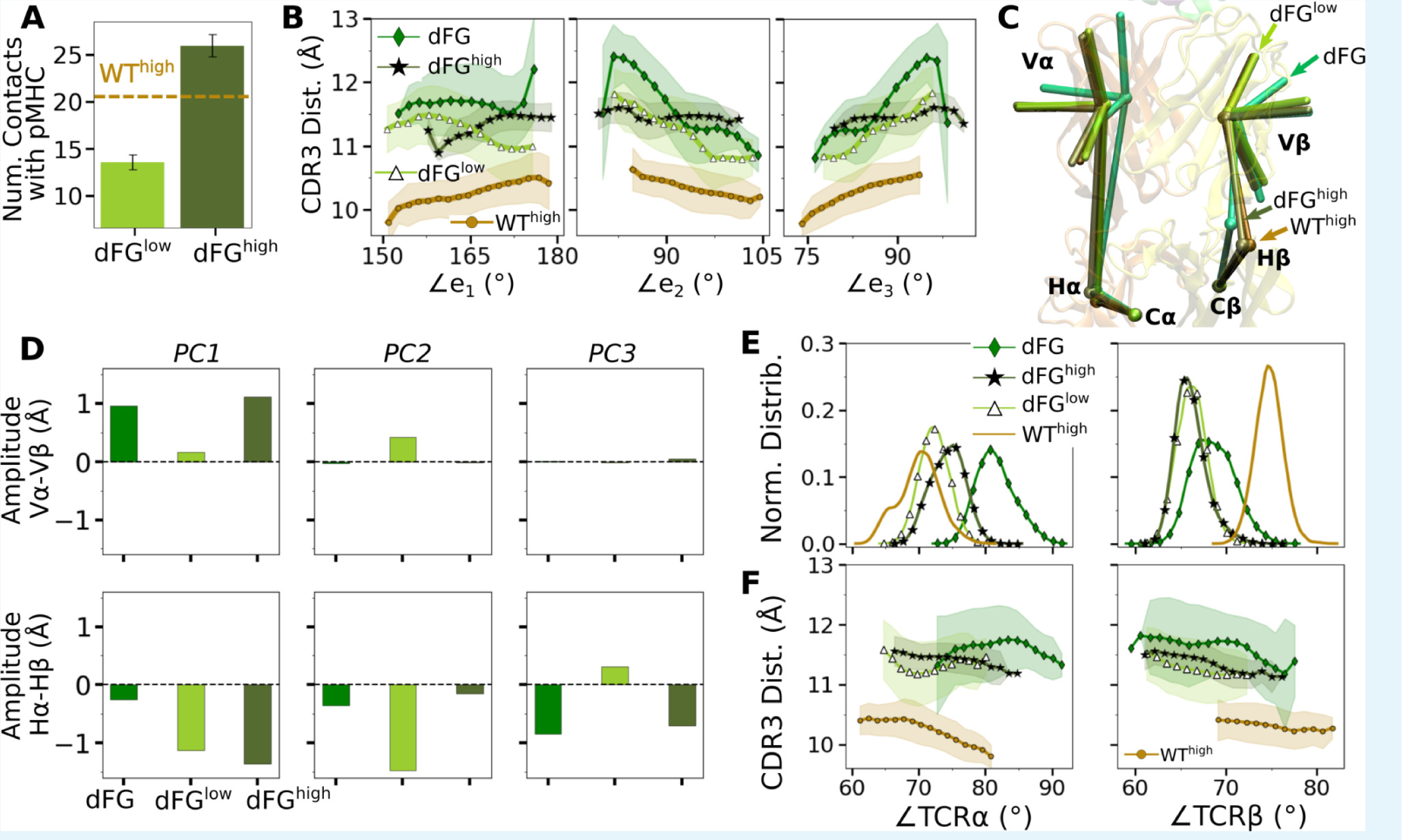
Effects of the C*β* FG-loop deletion. In all panels, the same criteria were used to measure values as for the WT systems in the corresponding figures. For comparison, respective data for WT^high^ are shown. (A) Number of contacts with pMHC (***Figure 2***A). (B) CDR3 distance vs. the three V*α*-V*β* triad arm angles (***Figure 3***D). (C) Average BOC of labeled complexes oriented to the C-module of WT^high^. The unliganded dFG has notably different average BOC (***Figure 6***C,D). (D) Differences in amplitudes between respective PC components of the *α* and *β* chains (***Figure 4***C). (E) Histogram of hinge angles and (F) CDR3 distance vs. hinge angles (***Figure 4***D,E).

**Appendix 1—figure 2.**
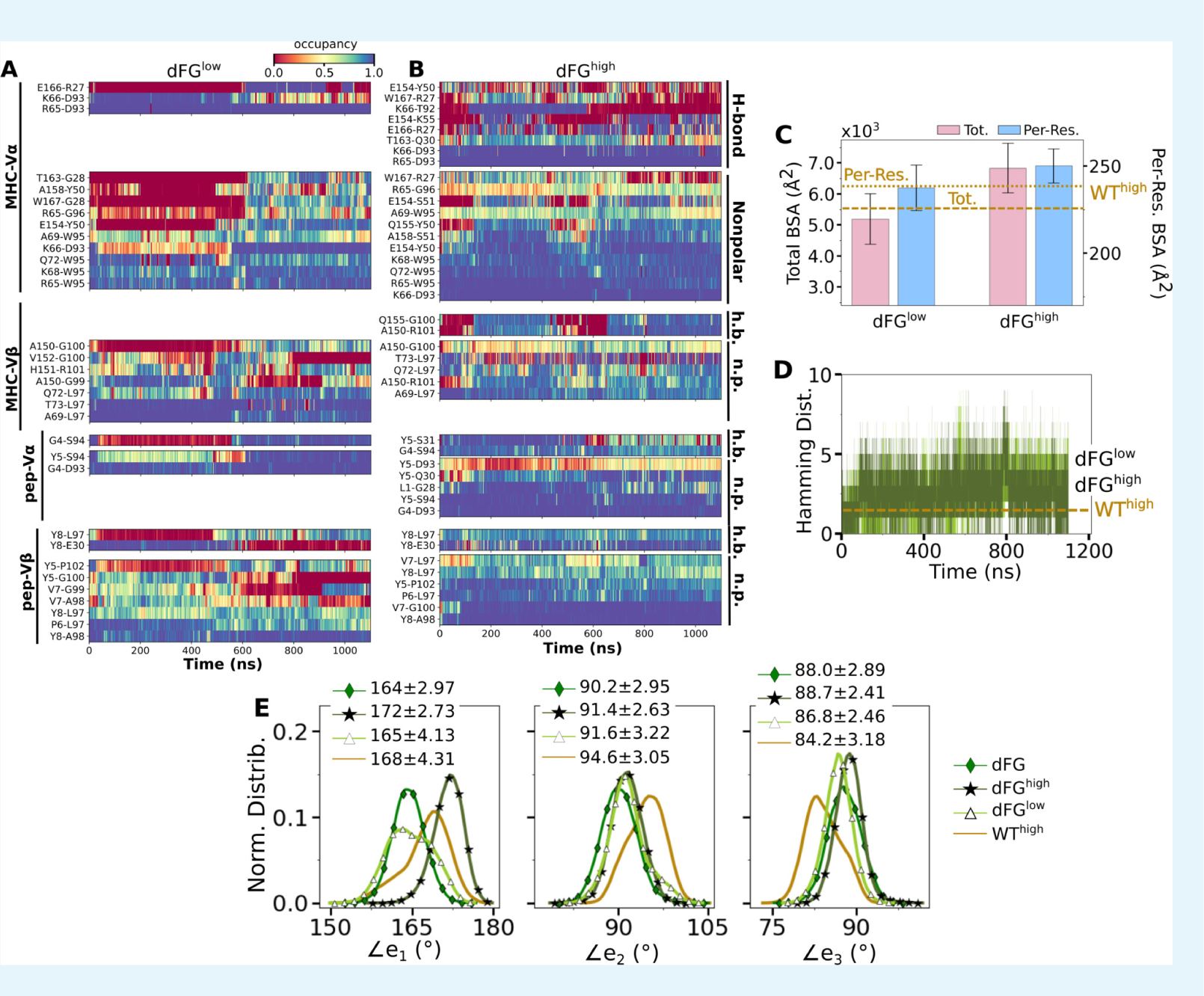
Effects of the C*β* FG-loop deletion on the interface with pMHC. The same criteria were used to plot as for the WT systems in the corresponding figures. (A.B) Contact
occupancy heat maps (Figure 2C,D). (C) BSA (Figure 2G). (D) Hamming distance (Figure 2B). (E) Distribution of V*α*-V*β*. angles. Numbers are avg±std in respective cases (Figure 3C). In (C.E), data for WT^high^ are shown as reference.

### Appendix 2

#### ***In silico*** peptide mutants mimic the behaviors of target systems

We tested whether behaviors of different systems are interchangeable by making point mutations on peptides *in silico*, for which WT and antagonists were used (Appendix 2–***Table 1***). For example, for ^Y8A^WT^high^ (Y8A under high load switched to WT), we took Y8A^high^ at the beginning of its production run and mutated A8 to Y8 (see *In silico* mutants in Methods). A main question is whether the *in silico* mutants attain behaviors of the switched systems during the finite simulation time. We found this to be the case although behaviors were not recapitulated perfectly. Compared to WT^high^, the number of contacts with pMHC became lower for ^WT^P6A (Appendix 2–***Figure 1***A). Note that ^WT^P6A^low^ had a higher force than ^WT^P6A^high^ (24.7 vs. 19.5 pN; Appendix 2–***Table 1***). This was because the low extension in the former case allowed the loose C-terminal strands (***Figure 1***A) to form extensive nonpolar contacts with the C-module, especially with C*β*. This effectively shortened the length of the complex, which led to a higher average force as the extension was kept the same (Selecting extensions). Among the original systems, Y5F^high^ had average force comparable to ^WT^P6A^low^ (23.7 pN; ***Table 1***), yet it had 1.8-fold more contacts, suggesting that the latter does behave like an antagonist (***Figure 5***A vs. Appendix 2–***Figure 1***A). For *in silico* WT, the number of contacts with pMHC was comparable to that of WT^high^ except for ^Y8A^WT^high^ (Appendix 2– ***Figure 1***A). In the original Y8A^high^, even though the number of contacts was at the level of WT^high^ (***Figure 5***A), the smaller size of A8 caused CDR3*β* to extend. The altered interface can be seen by the initial rapid increase in  (***Figure 5—figure Supplement 1***D). Mutating A8 to Y8 thereby forces the bulkier Y8 side chain to take an orientation different from that of WT. Thus, an *in silico* mutation of a residue to a comparable or smaller one is better tolerated than mutating to a bulkier one.

**Appendix 2—table 1.**
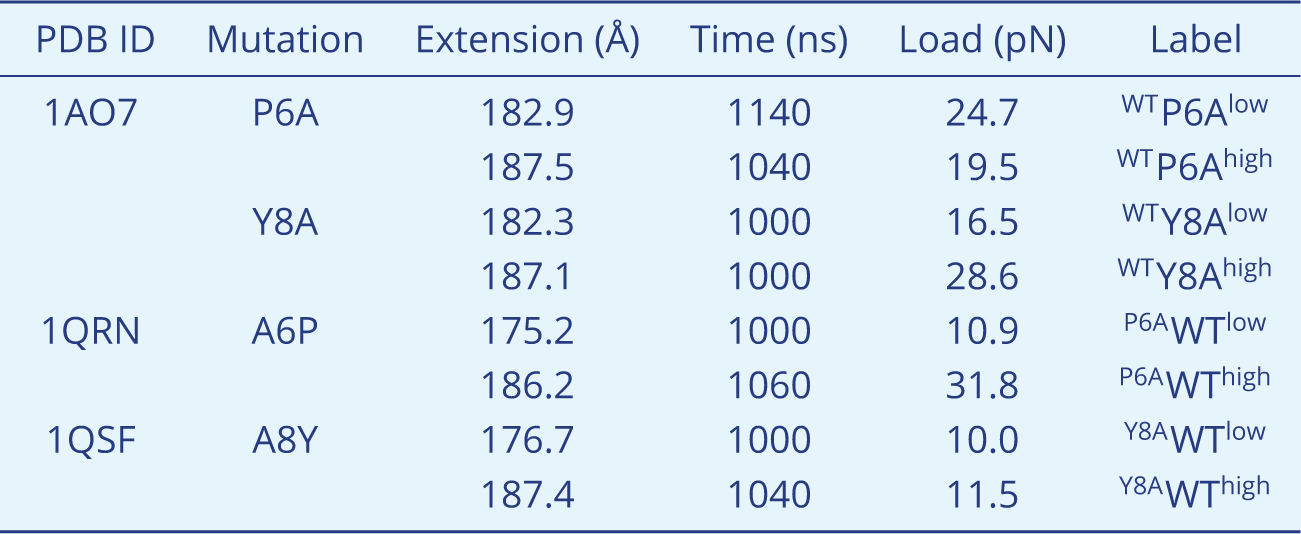
Simulations of TCR*αβ* with *in silico* mutations on the peptide. Load reported is average after 500 ns.

For ^WT^Y8A and ^P6A^WT, a higher load led to more contacts with pMHC (Appendix 2–***Figure 1***A). For ^WT^Y8A^high^, the number was comparable to WT^high^, which agrees with the case for the original Y8A^high^ (***Figure 5***A). The BSA profiles of *in silico* mutants also followed a trend similar to the number of contacts with pMHC, which was lower for antagonists and higher for WT (Appendix 2–***Figure 2***A). Differences in binding with pMHC can also be seen in the positional distribution of high-occupancy contacts, where ^P6A^WT had a relatively compact and evenly distributed contacts (Appendix 2–***Figure 2***B), although not as extensive as V*αβ*-pMHC or WT^high^ (***Figure 2***F).

Regarding the V*α*-V*β* interface, there were overall less contacts in the *in silico* antagonists than *in silico* WT (Appendix 2–***Figure 2***C). However, a higher load did not result in greater number of V*α*-V*β* contacts. The CDR3 distance was higher for *in silico* antagonists (Appendix 2–***Figure 1***B, top), while for *in silico* WT their range became narrower, to 11–12 Å which is similar to that for the modified agonist Y5F (***Figure 6***A vs. Appendix 2–***Figure 1***B, bottom row). The CDR3 distance also stabilized over time in all *in silico* WT, which was less so for *in silico* antagonists (Appendix 2–***Figure 2***D). However, similarly as the number of V*α*-V*β* contacts, there was no consistent load-dependence between the CDR3 distance and triad arm angles. On the other hand, there was a stronger load dependence in the average V-C BOC. The *in silico* antagonists that were built based on WT bent towards those of the corresponding antagonists, though the extent was not large (***Figure 6***D vs. Appendix 2–***Figure 1***C). The *in silico* WT in low load had average BOCs similar to those of the original antagonists whereas average BOCs of high-load *in silico* WT approached those of the actual WT (Appendix 2–***Figure 1***D). For ^Y8A^WT, this happened even though the forces experienced at the two extensions were only marginally different (10.0 vs. 11.5 pN; Appendix 2–***Table 1***). The V*α*-C*α* and V*β*-C*β* contacts were respectively lower for *in silico* WT than *in silico* antagonists, suggesting effects of the *in silico* mutations of the peptide did not propagate sufficiently to the whole TCR*αβ* during the simulation time (Appendix 2–***Figure 2***E). The lower number of V-C contacts in *in silico* WT would have made it easier to unbend under higher load or extension.

The above results suggest that the *in silico* mutants behave like the target system to varying extents. This is likely because the rearranged interface between the V-module and pMHC of the base system cannot immediately be adjusted upon *in silico* mutation in loaded states.

**Appendix 2—figure 1.**
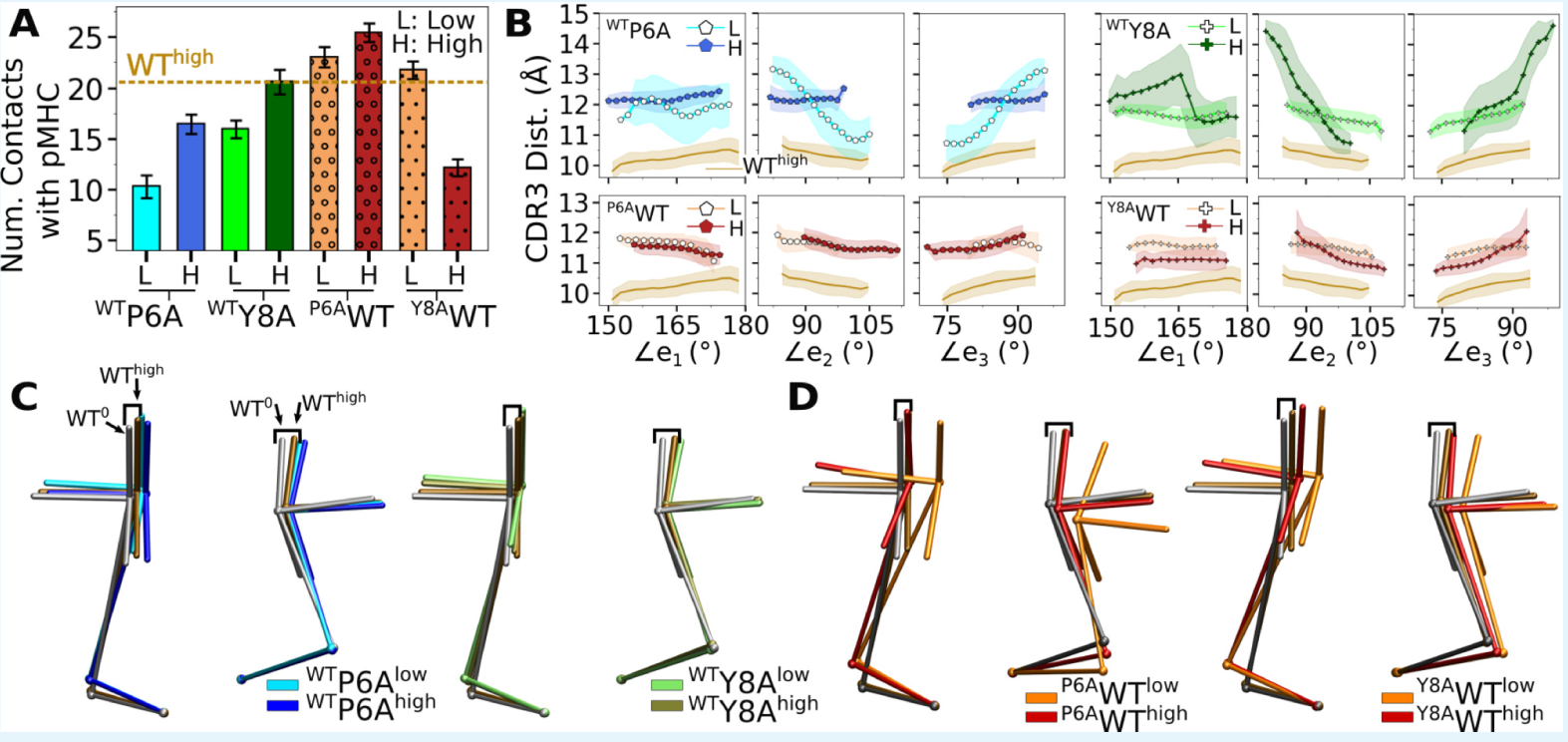
Simulations of *in silico* peptide mutants bound to A6. (A) Number of contacts with pMHC. Counts were made in the same way as in ***Figure 2***A. Dashed line: average value for WT^high^. Bars: std. (B) CDR3 distance vs. triad arm angles (***Figure 3***D and ***Figure 6***A,B). (C,D) Average BOCs of (C) *in silico* antagonists, and (D) *in silico* WT. Average BOCs of WT^0^ and WT^high^ are shown as reference, marked by angular brackets (***Figure 6***C,D).

**Appendix 2—figure 2.**
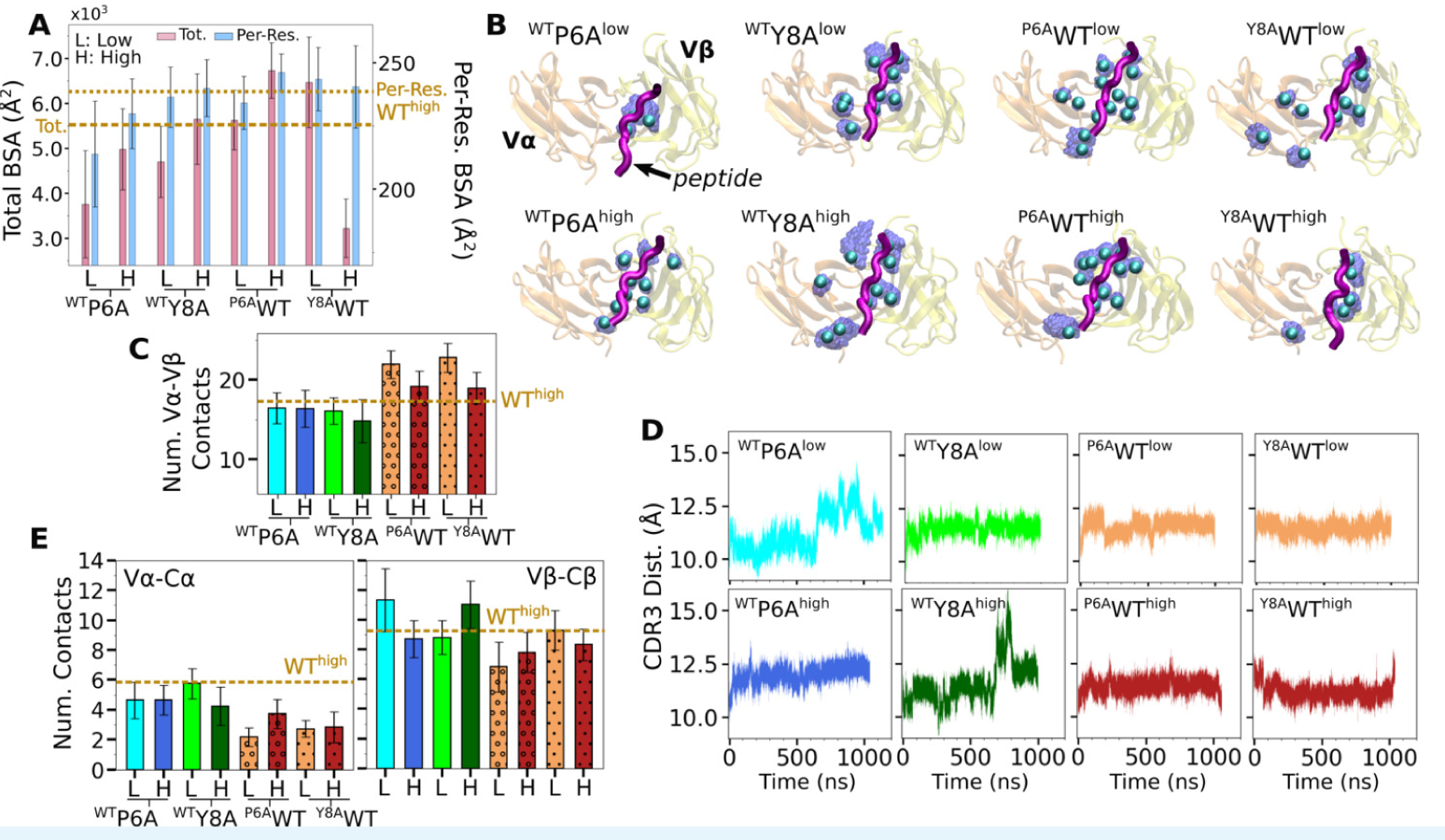
Behaviors of *in silico* mutants. (A) BSA (Figure 2G). (B) Positions of backbone C_*α*_ atoms of high contact occupancy residues (Figure 2F). (C) V_*α*_-V*β*. contact count (Figure 3A). (D) CDR3 distance trajectory (Figure 3.figure Supplement 2A.C). (E) V_*α*_-C*α*. and V*β*-C*β* contact counts (Figure 4-figure Supplement 1B,C).

---

**Figure 2—figure supplement 1.**
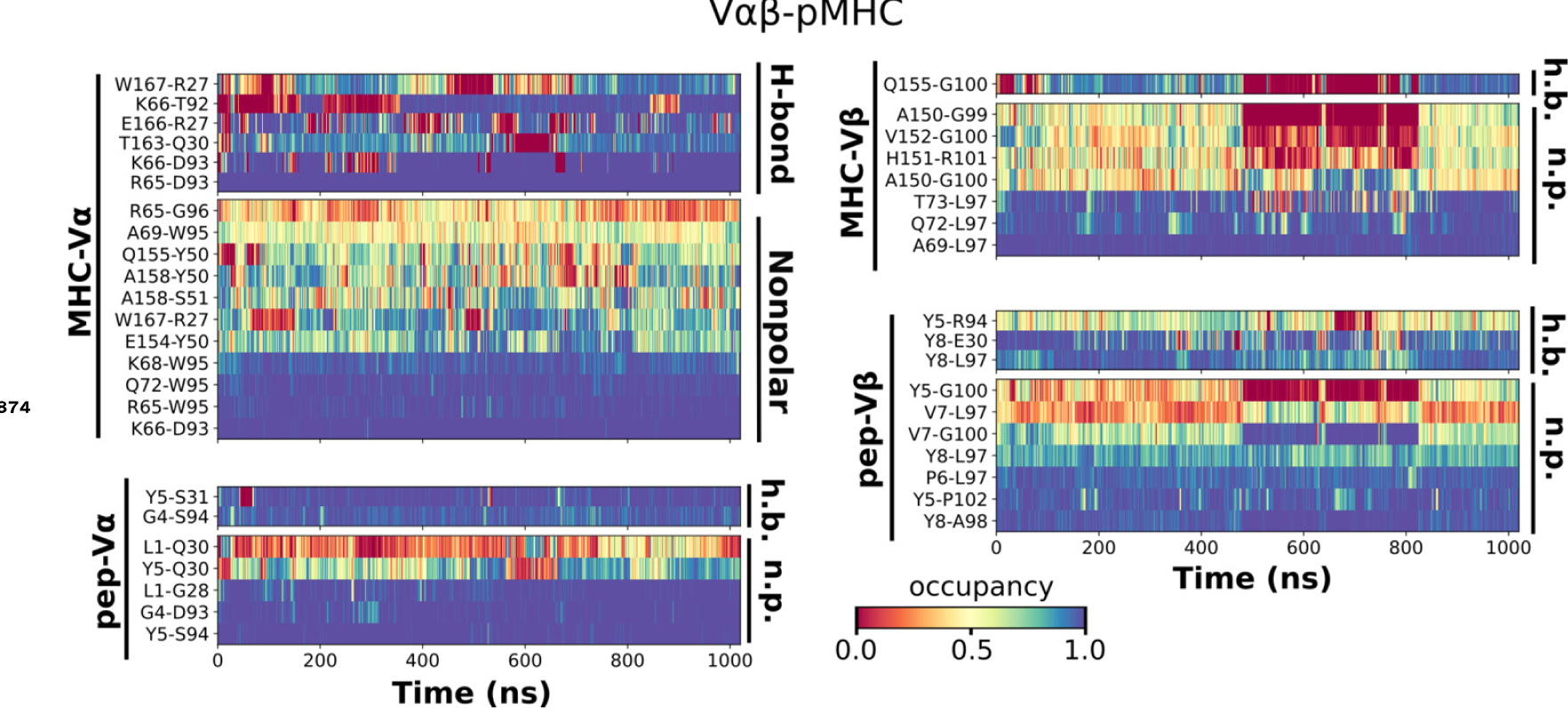
Contact occupancy heat maps for V*αβ*-pMHC. The same occupancy cutoffs as in Figure 2C–E were used.

**Figure 3—figure supplement 1.**
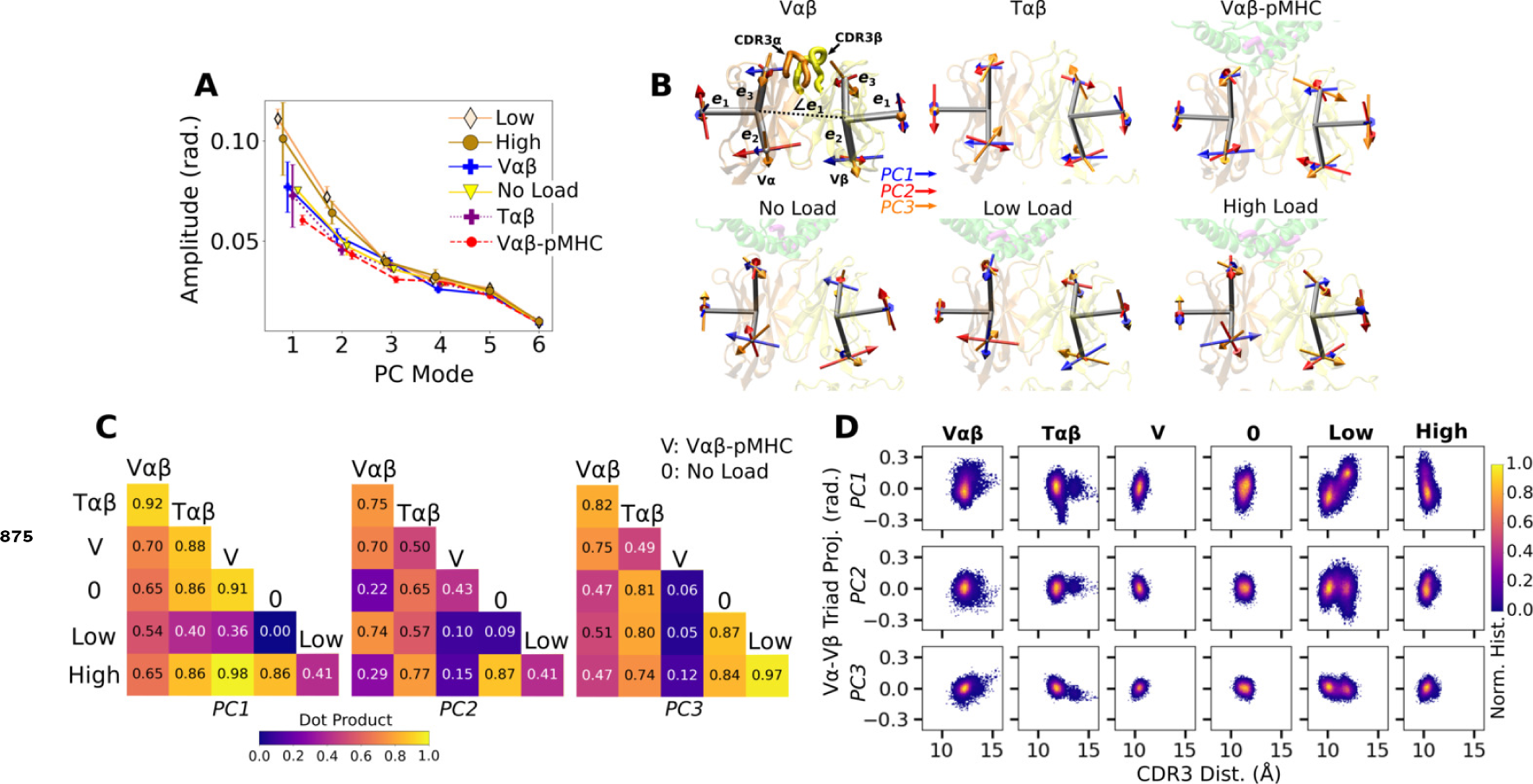
PCA of V*α*-V*β* motion. (A) PC amplitudes. Bars: std for PCA performed in 3 overlapping intervals from 500 ns to the end of simulation. (B) Direction of motion for the first three PC modes. (C) Absolute values of dot products between the unit PC vectors in listed systems. Values range from 0 (orthogonal) to 1.0 (identical PC directions). (D) 2-dimensional histograms of the projections of the V*α*-V*β* triads in each frame onto the first three PC directions versus the CDR3 distance.

**Figure 3—figure supplement 2.**
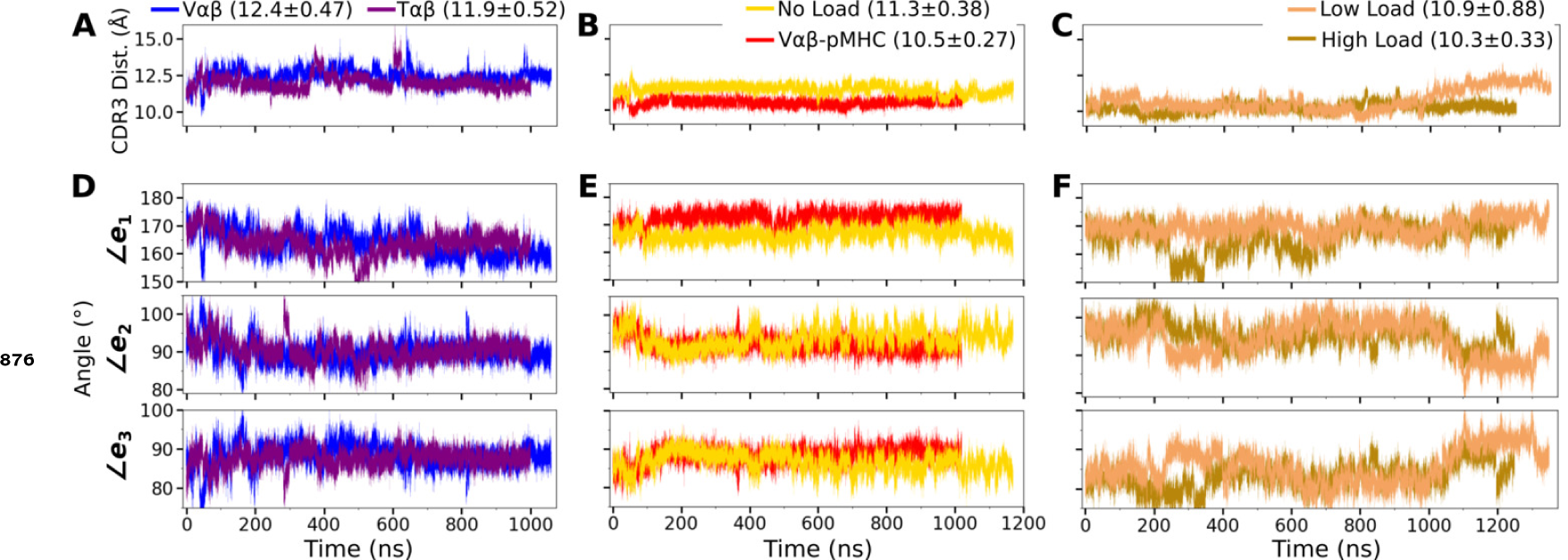
Trajectories of the V-module motion. (A–C) CDR3 distances and (D–F) triad angles. (A,D) V*αβ* and T*αβ*, (B,E) V*αβ*-pMHC and WT^0^, and (C,F) WT^low^ and WT^high^. Labels include average and standard deviation of the CDR3 distance after 500 ns.

**Figure 4—figure supplement 1.**
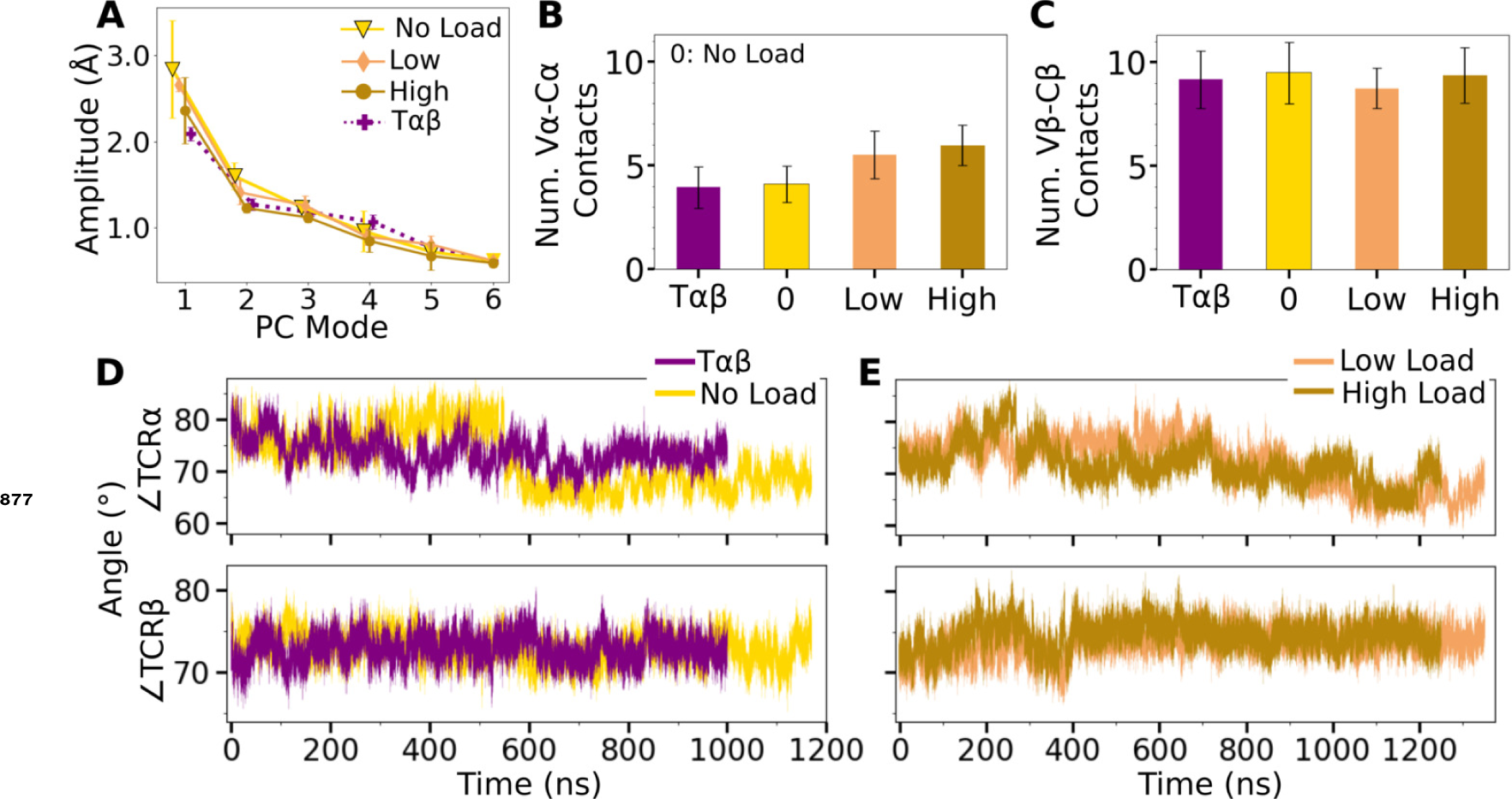
V-C PC amplitude and contacts. (A) Amplitude of the first six PCs. Bars: std for PCA performed in 3 overlapping intervals from 500 ns to the end of simulation. (B,C) Number of contacts with greater than 50% average occupancy and 80% maximum instantaneous occupancy for (B) V*α*-C*α* and (C) V*β*-C*β*. Bars: std. (D,E) Trajectories of hinge angles versus time.

**Figure 5—figure supplement 1.**
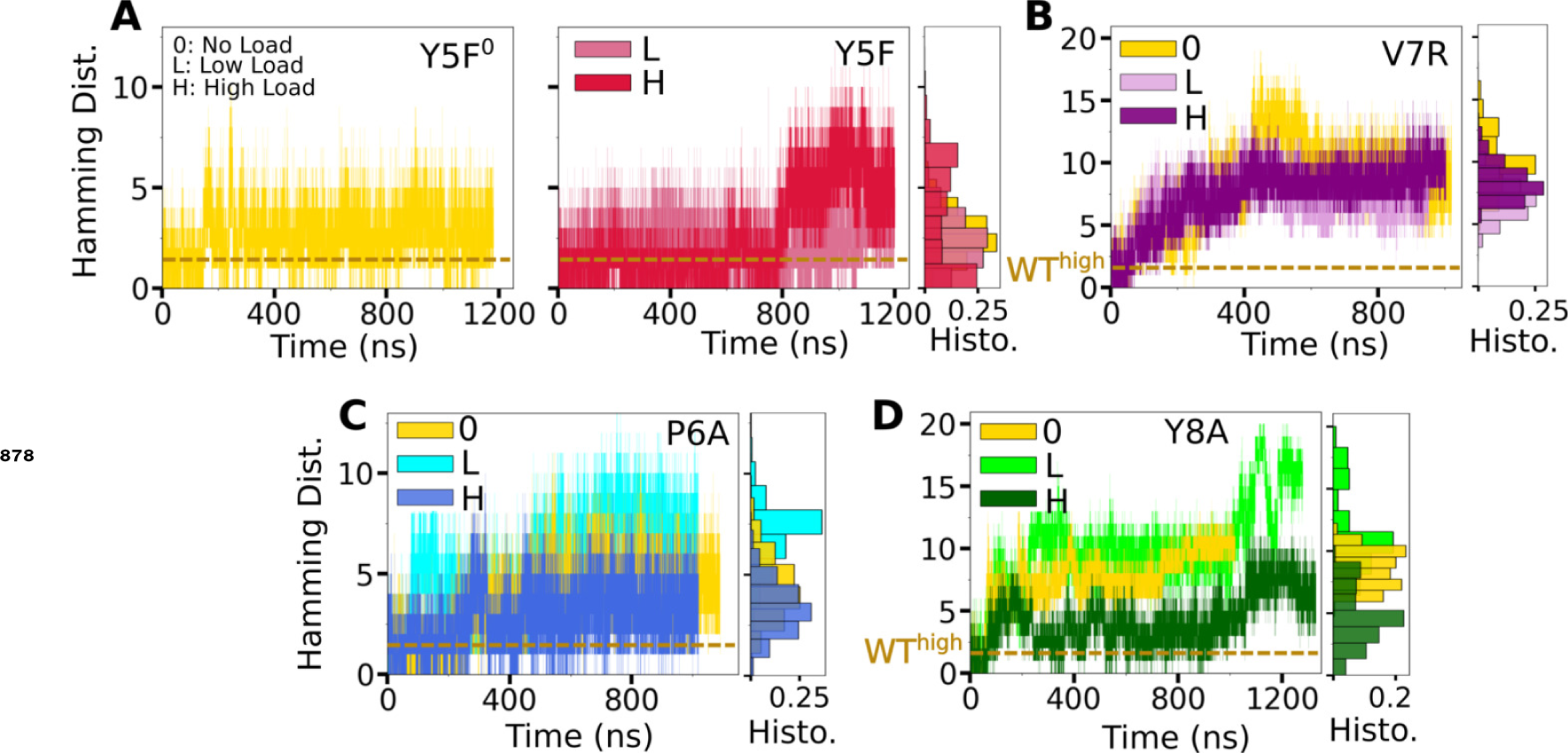
Trajectories of  for mutant complexes. (A,B) Modified agonists. (C,D) Antagonists. The same cutoff criteria were used to calculate initial contacts as in Figure 2B. Data after 500 ns were used for histograms on the right of each panel.

**Figure 5—figure supplement 2.**
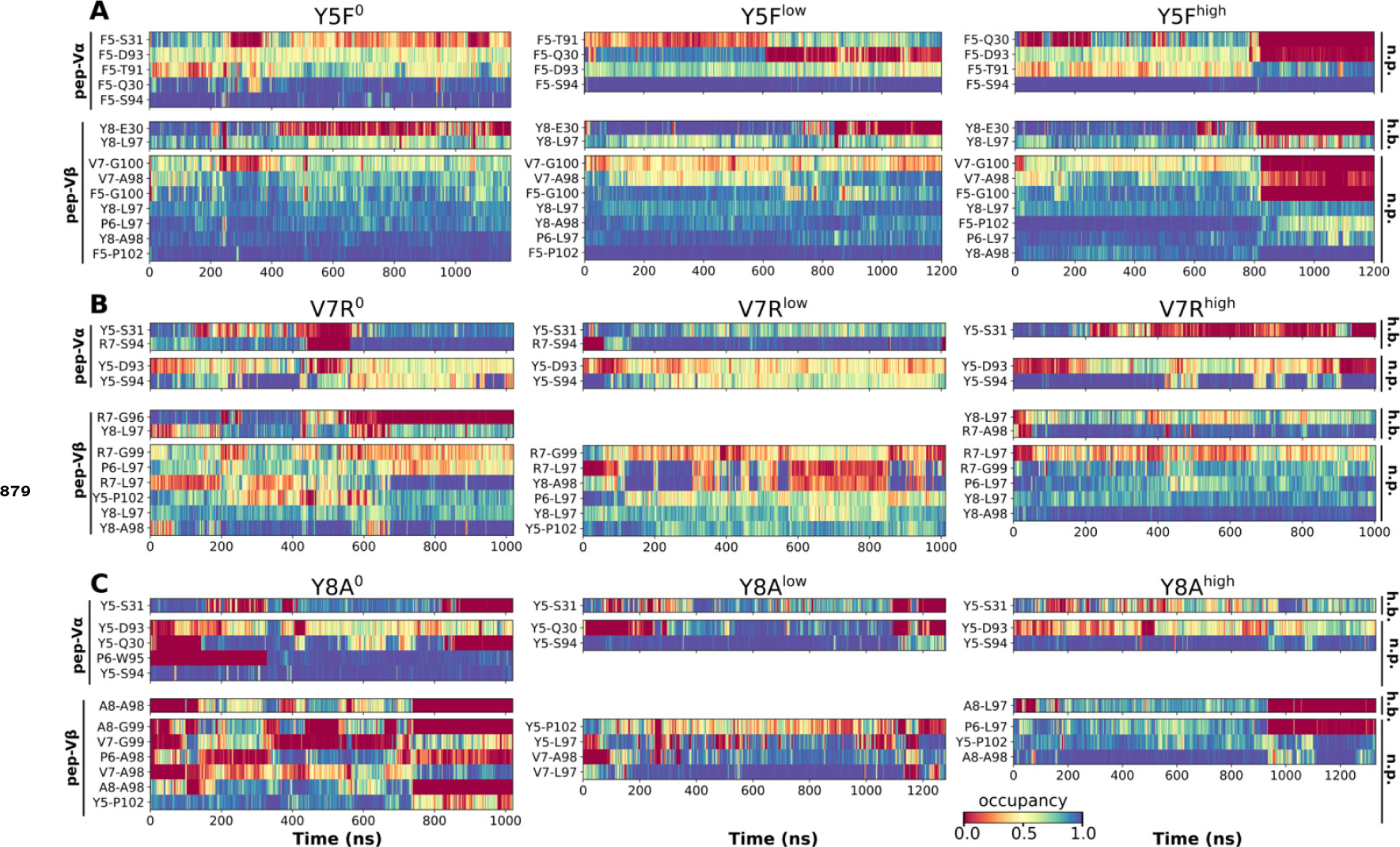
Contact occupancy heat maps for residues 5–8 of the mutant peptides. (A) Y5F, (B) V7R, and (C) Y8A. Corresponding heat maps for WT^high^ and P6A are in Figure 5C–F.

**Figure 5—figure supplement 3.**
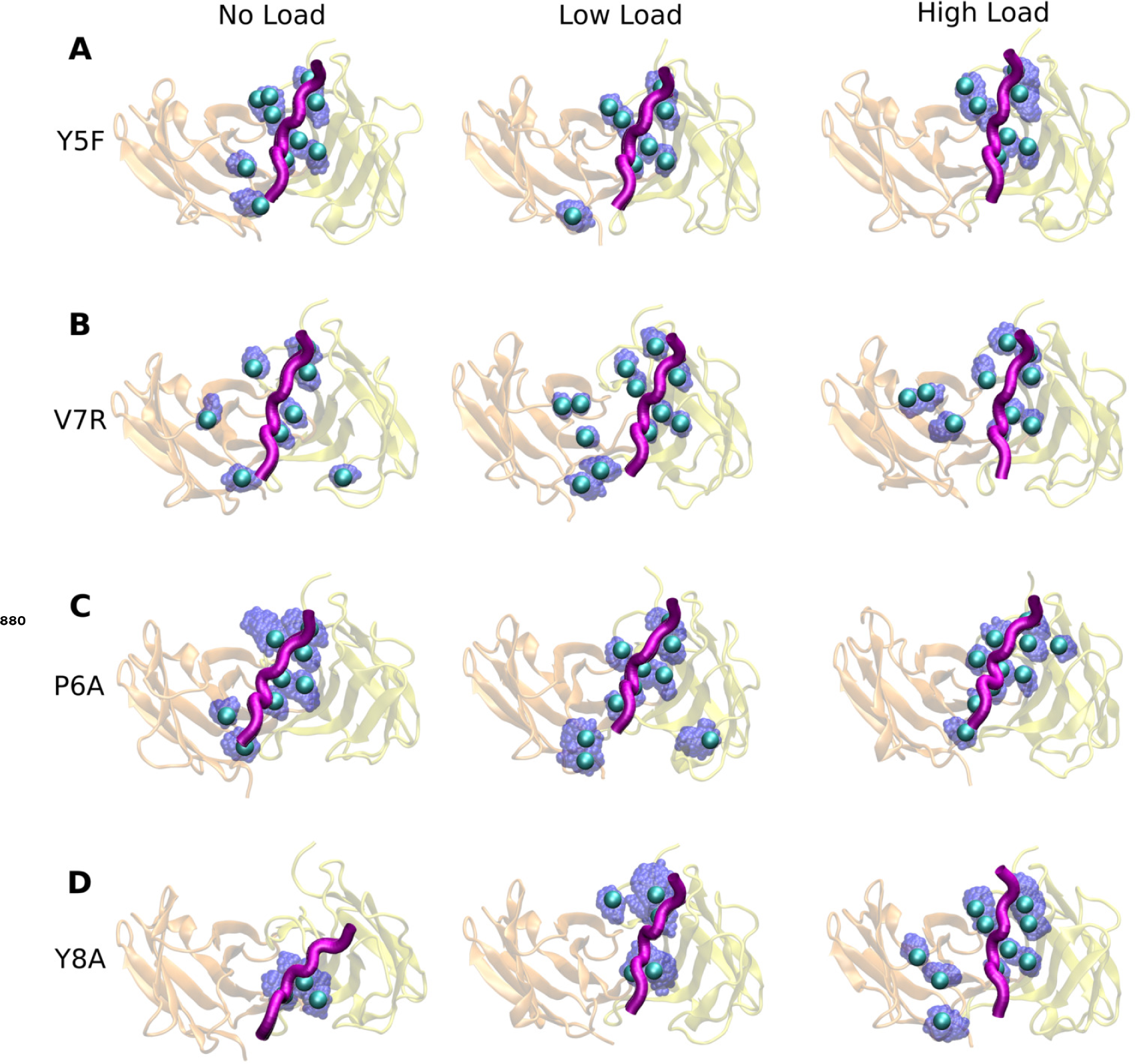
Locations of high-occupancy contacts with pMHC in mutant systems. (A) Y5F, (B) V7R, (C) P6A, and (D) Y8A. Compared to WT^high^ or V*αβ*-pMHC (Figure 2F), contacts are overall unevenly distributed or dispersed. The same occupancy cutoffs as in Figure 2F were used for selecting residues.

**Figure 6—figure supplement 1.**
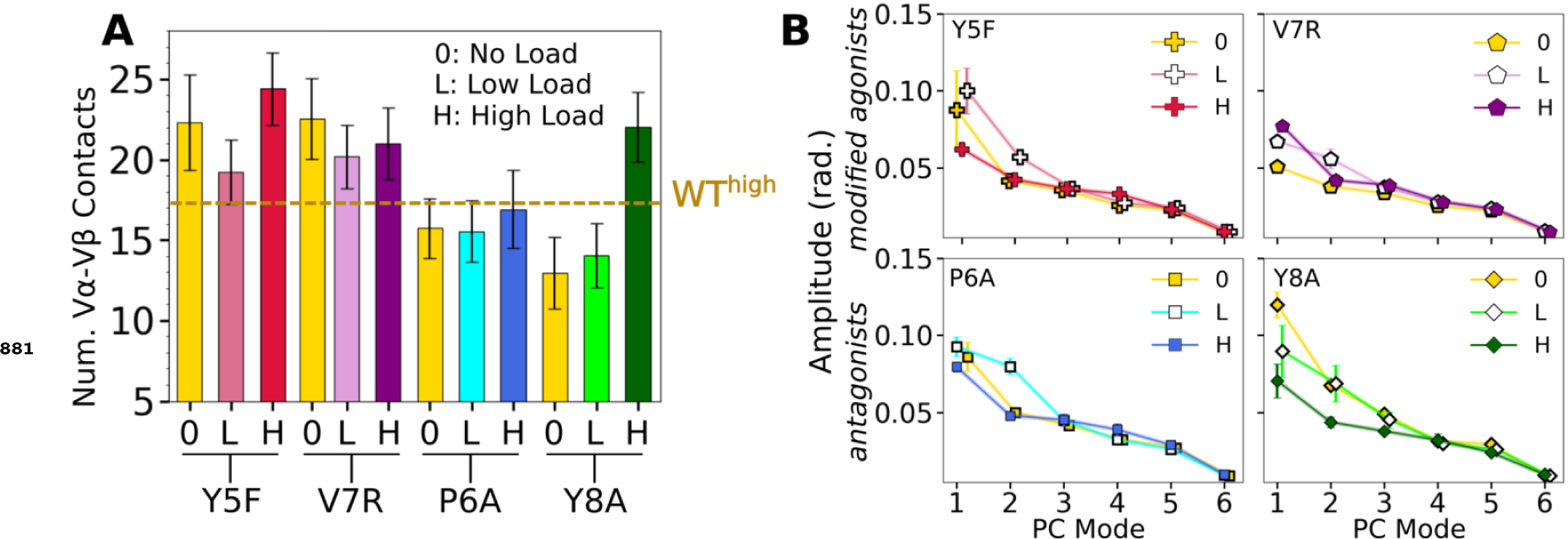
V*α*-V*β* motion of mutant systems. (A) Number of V*α*-V*β* contacts, counted in the same way as in Figure 3A. Dashed line is the average for WT^high^. (B) V*α*-V*β* PC amplitudes. Calculated the same way as in ***Figure 3—figure Supplement 1***A.

**Figure 6—figure supplement 2.**
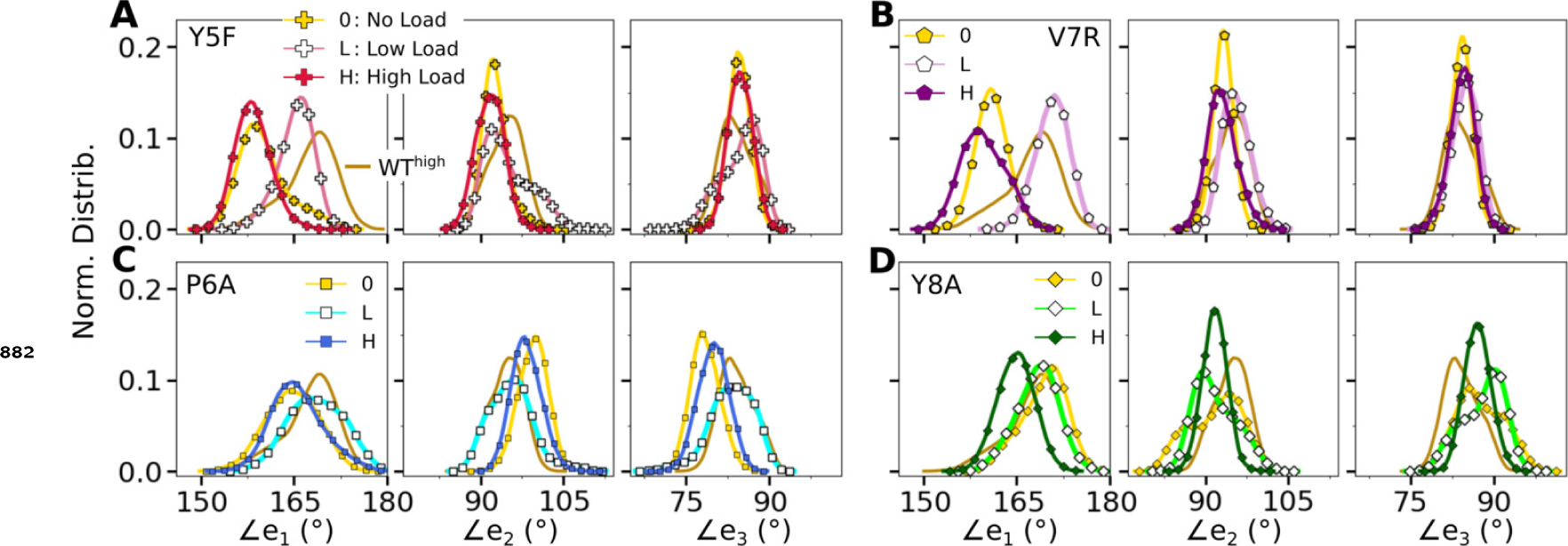
Distribution of triad arm angles in mutant systems. (A,B) Modified agonists and (C,D) antagonists. Respective plot for WT^high^ in Figure 3C is included in all panels (without markers) for comparison.

**Figure 6—figure supplement 3.**
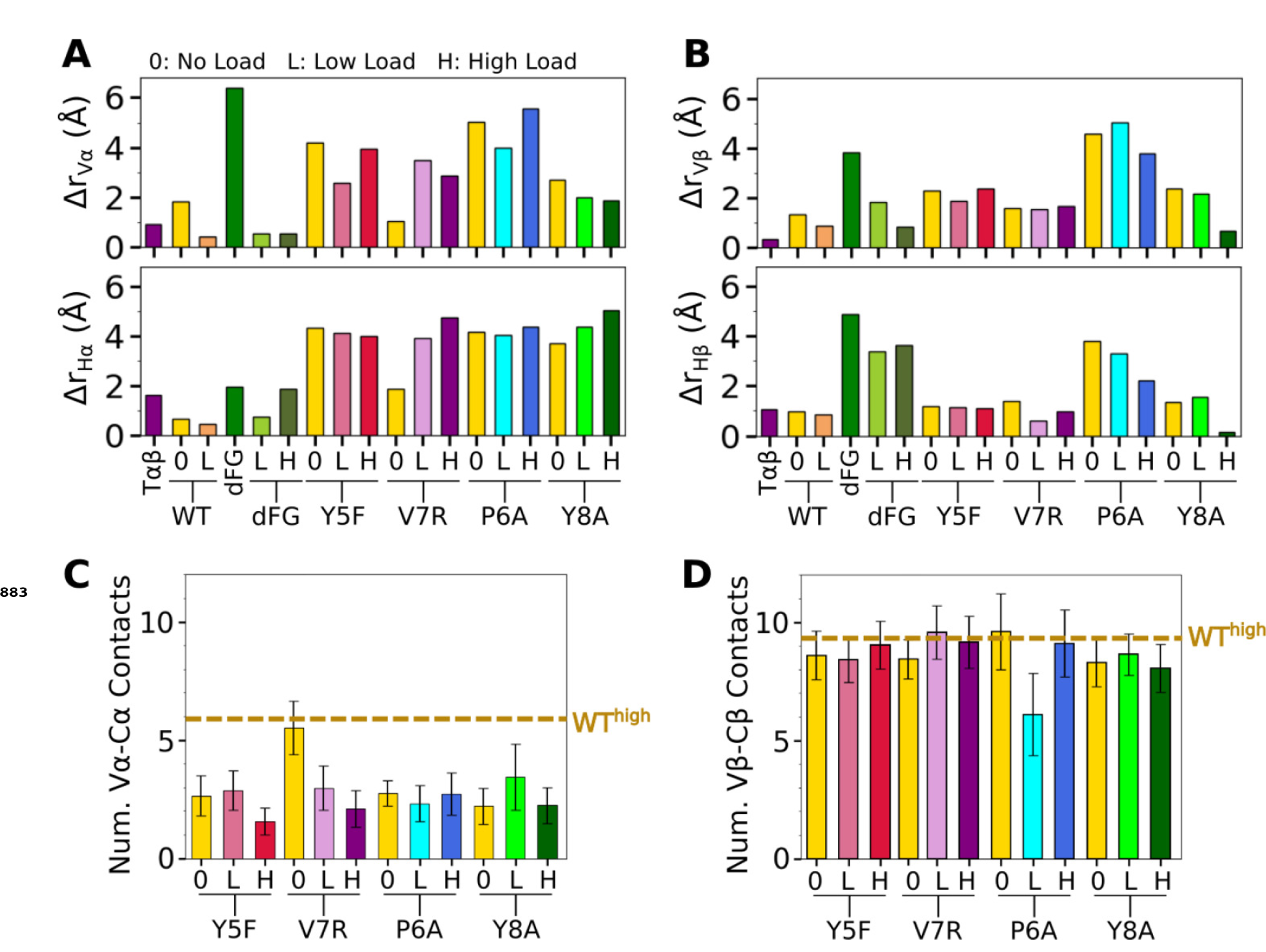
Comparison of mutant average V-C BOCs and interfaces with those of WT^high^. All BOCs are aligned to the C-module of WT^high^. (A,B) Distances of beads for (A) V*α* and H*α*, and (B) V*β* and H*β* from those of WT^high^, revealing the extent of deformation. (C,D) Number of V-C contacts for each chain. Dashed line denotes respective value for WT^high^ in ***Figure 4—figure Supplement 1***B,C.

**Figure 6—figure supplement 4.**
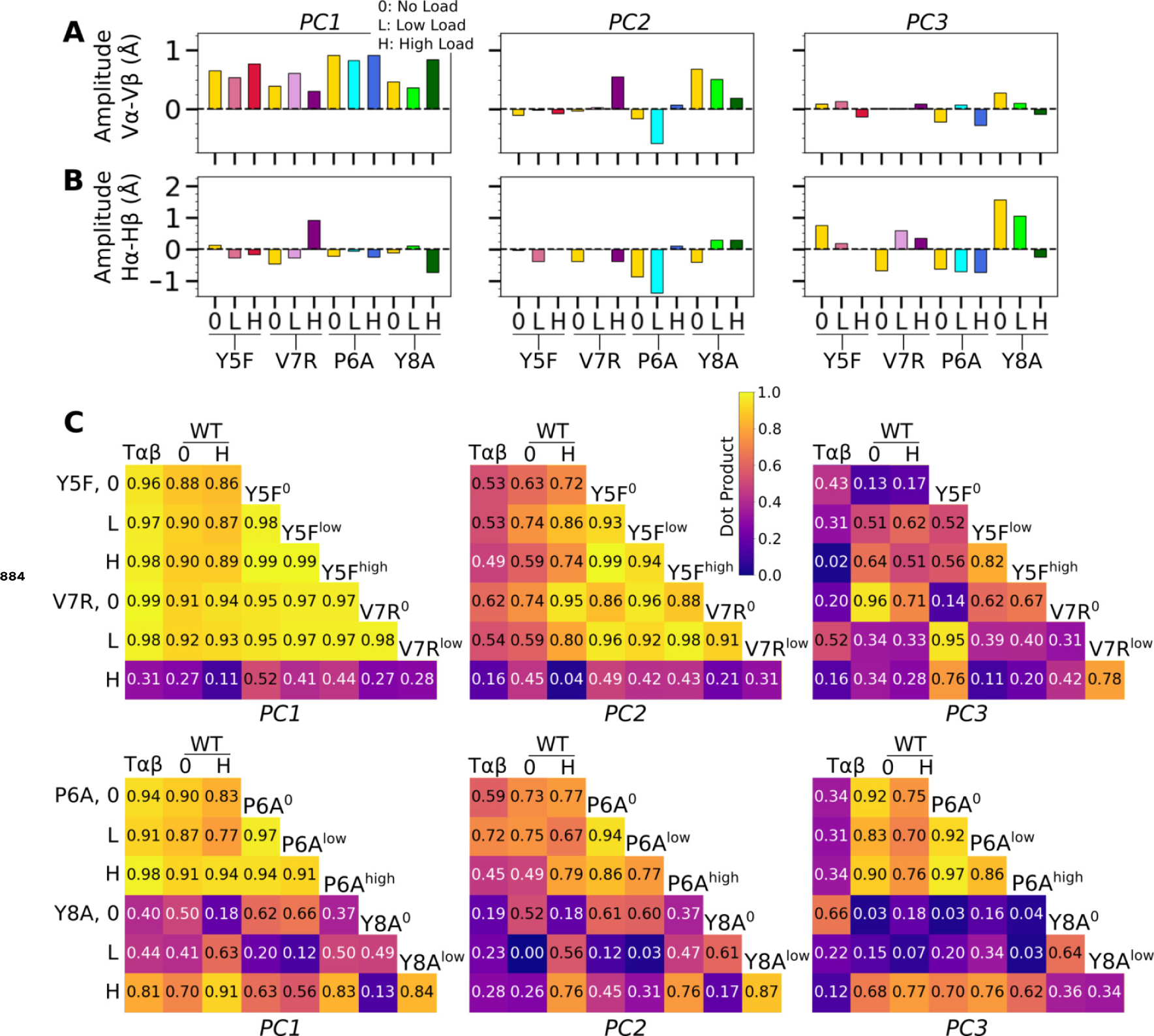
V-C motion of mutants. (A,B) Differences in PC amplitude between BOC PC of (A) V*α* vs. V*β* and (B) H*α* vs. H*β*. Compare with Figure 4C for WT systems. (C) Dot products between BOC PC vectors for the listed systems.

**Figure 7—figure supplement 1.**
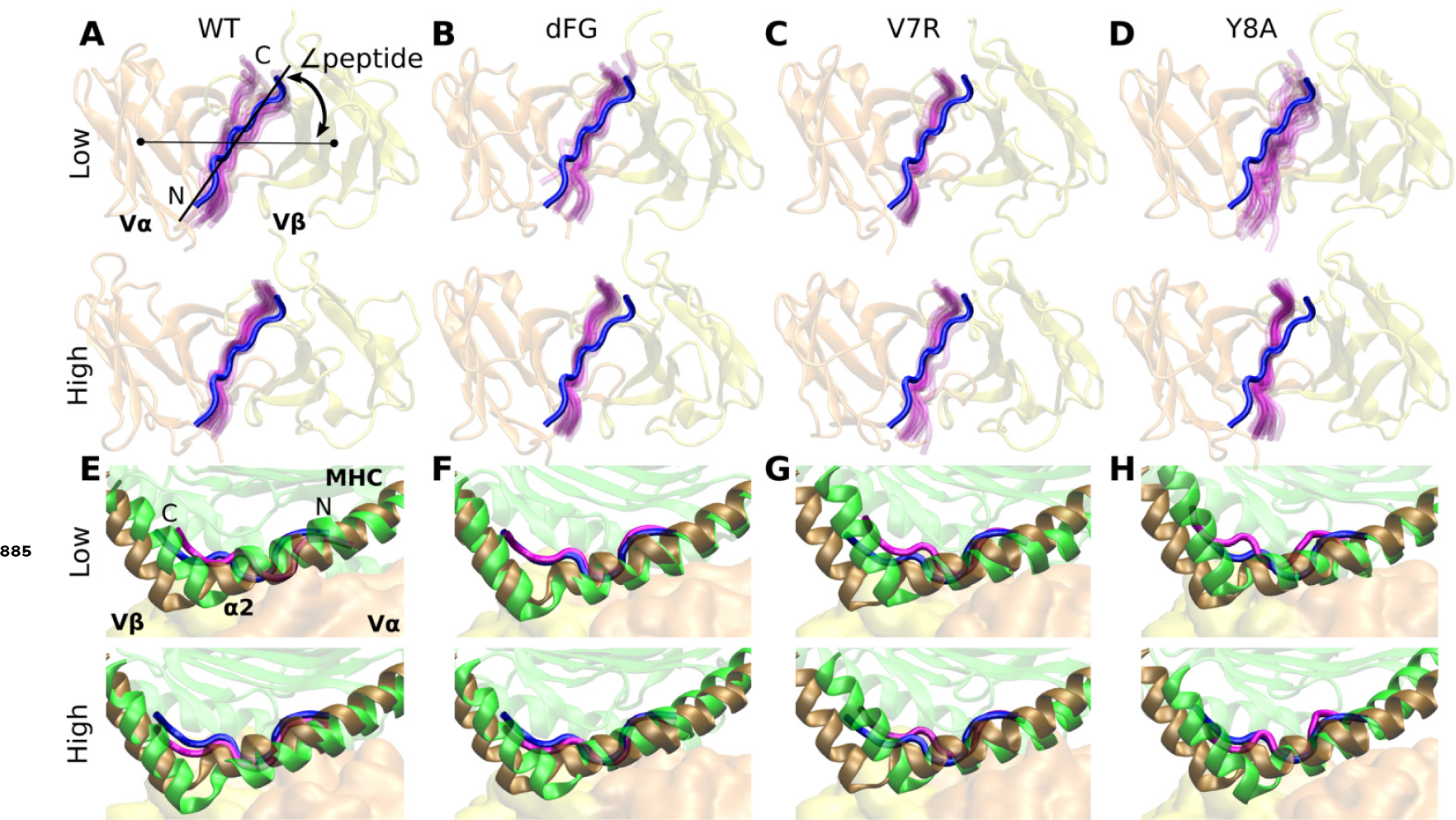
Motion at the interface related to ∠peptide. (A–D) View of interface from the top of the V-module. The peptide from the crystal structure (blue) of each respective system is overlaid on frames of the peptide during simulation (magenta) rendered every 50-ns from 500 ns to the end. (E–H) Positional shift of pMHC. Side view showing MHC *α*2 helix (brown) and peptide (blue) from the crystal structure overlaid with MHC *α*2 helix (green) and peptide (magenta) of the last rendered frame from panels A–D.

